# Senescence-directed nanotherapy ameliorates fibrosis and overcomes immune exclusion in cancer

**DOI:** 10.64898/2026.03.02.706451

**Authors:** Clemens Hinterleitner, Valentin J. A. Barthet, Hailey V. Goldberg, Kristen C. Vogt, Ana Marie Perea, Logan R. Hillger, Stephen Ruiz, Domhnall McHugh, Yu-Jui Ho, Almudena Chaves-Perez, Maria Skamagki, Sara Flowers, Hannah C. Styers, Natasha Rekhtman, Xueqian Zhuang, Gabriel Dessotti Barretto, Xiang Li, Jadae T. Watson, Wei Luan, Janelle Simon, Tuomas Tammela, Rui Gardner, Charles M. Rudin, Paul B. Romesser, Matthew J. Bott, Aveline Filliol, Daniel A. Heller, Scott W. Lowe

**Affiliations:** Cancer Biology and Genetics Program, Sloan Kettering Institute, Memorial Sloan Kettering Cancer Center, New York, NY, 10065, USA; Graduate School of Medical Sciences, Weill Cornell Medicine, New York, New York 10065, USA; Molecular Pharmacology Program, Memorial Sloan Kettering Cancer Center, New York, NY 10065, USA; Tri-Institutional PhD Program in Chemical Biology, Memorial Sloan Kettering Cancer Center, New York, NY 10065, USA; Cancer Engineering Program, Gerstner Sloan Kettering School for Biomedical Sciences, Memorial Sloan Kettering Cancer Center, New York, NY 10065, USA; Department of Surgery, Memorial Sloan Kettering Cancer Center, New York, New York 10065, USA; Department of Radiation Oncology, Memorial Sloan Kettering Cancer Center, New York, NY, 10065, USA; Department of Pathology, Memorial Sloan Kettering Cancer Center, New York, NY, 10065, USA; Flow Cytometry Core Facility, Memorial Sloan Kettering Cancer Center, New York, NY, 10065 USA; Department of Medicine, Memorial Sloan Kettering Cancer Center, New York, NY 10065, USA; Early Drug Development Service, Department of Medicine, Memorial Sloan Kettering Cancer Center, New York, NY, 10065, USA; Howard Hughes Medical Institute, Chevy Chase, MD 20815, USA

## Abstract

Fibrotic remodeling of tissues and tumors establishes immunosuppressive microenvironments that drive organ dysfunction and, in cancer, limit response to immunotherapy. Senescent-like cells are conserved drivers of fibrosis and therapeutic targets, yet their functional heterogeneity complicates therapeutic intervention. Here, we show that P-selectin is expressed by a subset of senescent-like cells in fibrotic tissues and tumors. Leveraging fucoidan-based nanoparticles that bind P-selectin, we developed senescence-modulating nanoparticles (SMNPs) to selectively target these disease-associated states. SMNPs exerted potent antifibrotic and immunomodulatory effects while improving therapeutic index. Mechanistically, we identified a pathogenic, immunosuppressive macrophage population as a functional target in vivo. In fibrotic tumors, niche remodeling restored immune infiltration and sensitized tumors to immune-checkpoint-based therapies. These findings establish SMNPs as a generalizable strategy to target pathogenic senescent cell subsets across fibrosis and cancer.

## Introduction

Chronic inflammation of the liver and lung frequently culminates in fibrosis, progressive organ dysfunction, and premature mortality, yet disease-modifying therapies remain limited (*1–4*). Fibrosis is marked by the accumulation of activated stromal and immune cells, excessive extracellular matrix deposition, vascular remodeling, and distortion of tissue architecture, collectively reshaping local cellular ecosystems (*5–7*). Additionally, fibrotic environments are potently immune suppressive, form barriers to immune surveillance that constrain host defense and limit the efficacy of immunotherapies in cancer (*8–11*). These observations imply that chronic inflammation, fibrosis, immune dysfunction, and cancer progression may reflect convergent biological programs rather than distinct pathological entities. Accordingly, identifying and targeting the cellular states that sustain fibrotic, immune-suppressive niches represents a shared challenge across chronic inflammatory disease and oncology.

One cellular process implicated in fibrosis is cellular senescence, a stress response program engaged by diverse forms of cellular and tissue injury (*12–16*). Senescence is associated with stable proliferative arrest and broad rewiring of cell physiology, including activation of tissue remodeling programs that alter how cells signal and sense their microenvironment (*17, 18*). While the presence of cells exhibiting senescence features can support tissue repair following acute injury, their persistence in chronically damaged tissues sustains inflammation, matrix remodeling, and architectural distortion, thereby accelerating fibrotic progression and organ decline (*17, 19–21*). In cancer, senescent-like cells accumulate within the tumor microenvironment (TME), where they are linked to stromal remodeling, altered myeloid cell composition, and immune suppression (*22–29*). Together, these observations position senescent cells as conserved drivers of fibrotic remodeling, immune dysfunction, and tissue decline across organs and disease contexts.

Pharmacologic strategies to eliminate or modulate senescent cells show promise in preclinical models (*30–35*), yet translation has been limited by two fundamental challenges (*16*). First, while senescence was defined in vitro, senescence-like states in tissues are heterogeneous, varying across cell types, tissue, and association with distinct functional consequences (*14, 36*). Second, existing senotherapeutics lack cellular precision, leading to predictable on-target toxicity in normal or reparative cell populations (*30, 34, 37–39*). These limitations underscore the need for strategies that selectively target pathogenic senescent states while sparing beneficial counterparts, a goal that requires improved resolution of senescent-state heterogeneity in vivo and its relationship to tissue pathology.

Motivated by our finding that P-selectin (*SELP*) is upregulated on cells with senescent features, we reasoned that this molecule could provide a tractable entry point for selective therapeutic targeting. P-selectin is induced on activated endothelium in inflammatory and fibrotic settings, where it mediates leukocyte recruitment and transcytosis via PSGL-1 and related glycosylated ligands (*40–45*). Fucoidan, a sulfated polysaccharide and high-affinity P-selectin ligand, can be formulated into nanoparticles that encapsulate small-molecule drugs, enabling P-selectin–directed delivery of therapeutic payloads (*46–48*). We therefore hypothesized that P-selectin could function both as a conduit for tissue-selective drug delivery and a molecular handle to preferentially access senescent-like, pathology-driving cell states within fibrotic niches. In this study, we define the landscape of P-selectin expression across senescent-like states in fibrotic tissues and evaluate the therapeutic potential of P-selectin–directed senescence-modulatory nanoparticles (SMNPs) in preclinical models of fibrosis and cancer.

## Results

### P-selectin marks senescent-like cells in fibrotic tissues

Given emerging links among P-selectin biology, senescence, and fibrosis (*42–45*), together with the potential to exploit P-selectin-dependent pathways for targeted drug delivery, we sought to systematically define the relationship between P-selectin expression and senescent cell states in fibrotic disease. We analyzed 58 human liver and lung fibrosis specimens, alongside healthy controls, using conventional immunohistochemistry and multiplexed immunofluorescence (mIF) panels incorporating canonical senescence markers. Across both organs, P-selectin expression was significantly elevated in fibrotic tissues compared to non-fibrotic counterparts (Fig. 1A–C and fig. S1 A-B). Within these tissues, a subset of P-selectin-positive cells exhibited multiple hallmark features of cellular senescence, including expression of the cell-cycle inhibitor p21, absence of the proliferation marker Ki67, and reduced expression of the nuclear lamina protein LaminB1 (Fig. 1A-C and fig. S1C). We refer to such states as “senescent-like” to acknowledge the heterogeneity of senescence programs and their incompletely defined nature in vivo (*16, 49, 50*).

**Fig. 1:**
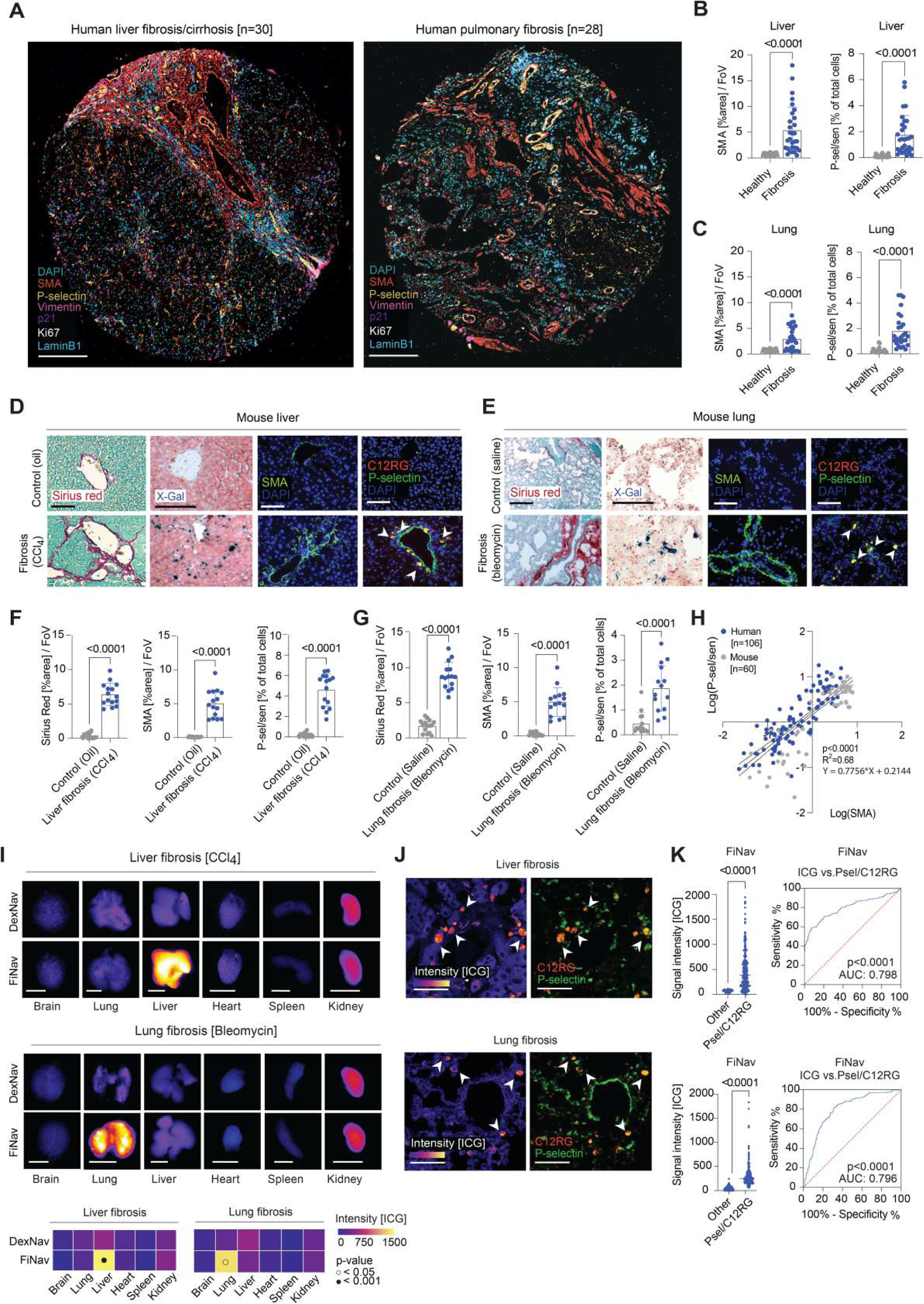
P-selectin-expressing senescent-like cells in fibrotic tissues are selectively targeted by fucoidan nanoparticles. (**A-C**) mIF analysis of tissue microarrays of human liver (healthy, n=21; fibrotic/cirrhotic, n=30 cases) and lung (healthy, n=21; fibrotic, n=28 cases). The figure shows representative images of fibrotic organs (A) and quantification of SMA^+^ activated fibroblasts and P-selectin^+^ senescent cells (defined as p21^+^, Ki67^-^, LaminB1^low^) (B-C). Scale bars, 500 μm. (**D-G**) Collagen deposition (Sirius Red), senescence-associated-β-galactosidase activity (X-Gal), SMA, and P-selectin expression in senescent cells (defined as C12RG^+^) in murine models of healthy liver (oil treated), liver fibrosis (6 weeks after start of CCl_4_ treatment) (D) and healthy lung (saline) and lung fibrosis (28 days after bleomycin treatment) (E). The figure shows representative images (D,E) and quantification (F,G). Data represent 13-15 mice per group from three independent experiments. Scale bars, 100 μm. (**H**) Pearson correlation analysis between P-selectin^+^ senescent cells and activated fibroblasts in human and murine fibrotic and healthy liver and lung tissues. (**I**) Near-infrared fluorescence imaging of indocyanine green-positive (ICG^+^) nanoparticles (FiNav and DexNav) across tissues in murine models of liver fibrosis (top) and lung fibrosis (middle) at 12h post nanoparticle injection. (Bottom) Quantification of ICG^+^ nanoparticle enrichment in each tissue (3 mice/group; one experiment). Scale bars, 500 μm. (**J**) Representative fluorescent images of FiNav enrichment (ICG^+^) in P-selectin^+^ senescent cells (C12RG^+^) of murine fibrotic liver (top) and lung (bottom). Scale bars, 50 μm. (**K**) Quantification of ICG in P-selectin^+^ senescent cells vs all other cell populations (3 mice per group, left) and receiver operating characteristic (ROC) curve of ICG specificity for P-selectin^+^ senescent cells (3 mice per group, right). Each dot represents one patient or one mouse (B-C; F-G, respectively) or one cell (K). Data are presented as mean ± s.e.m. Statistical tests were Mann Whitney (B-C, F-G, K), Welch’s t test (F, G), simple linear regression (H), one-way ANOVA with Sidak’s multiple comparisons (I), and ROC analysis (K).

We next asked whether the presence of P-selectin-positive senescent-like cells was conserved in murine fibrosis models, where senescent cells have been implicated as drivers of tissue pathology (*50, 51*). In CCl_4_- and bleomycin-induced models of liver and lung fibrosis (*52, 53*), we observed increased collagen deposition and an increase in the fraction of cells expressing senescence-associated beta-galactosidase (SA-β-gal), a canonical senescence marker (*54, 55*) (Fig. 1D-G). Consistent with human tissues, P-selectin expression was markedly increased in fibrotic murine livers and lungs across both species and organs. P-selectin expression was enriched in regions densely staining for α-smooth muscle actin (α-SMA), a marker of activated fibroblasts involved in extracellular matrix (ECM) deposition (*56, 57*) (Fig. 1D-G). Notably, a subset of P-selectin-positive cells also exhibited SA-β-gal activity, detected using the fluorescent substrate C12FDG (Fig. 1D-F). Collectively, these data establish that P-selectin marks a subset of senescent-like cells that accumulate in fibrotic environments, revealing a conserved association between P-selectin expression, senescence-associated features, and fibrosis across organs and species (Fig. 1H).

### SMNPs selectively target senescent-like cells in fibrotic organs

Having established that P-selectin is upregulated on senescent-like cells in fibrotic tissues, we next asked whether this feature could be exploited for targeted drug delivery. Leveraging our previously described fucoidan (Fi)-based, P-selectin–targeting nanoparticles (*48*), we engineered senescence-modulating nanoparticles (SMNPs) harboring agents known to impact senescence-associated phenotypes. One SMNP type encapsulated the senolytic agent Navitoclax, which targets BCL-2 family anti-apoptotic proteins induced in senescent states (*58, 59*), generating FiNav. The second SMNP type encapsulated the senomorphic agent dBET6, a BRD4-targeting proteolysis targeting chimera (PROTAC) that disrupts enhancer function and suppresses expression of SASP genes (*60*), generating FidBET6 (*47*). Nanoparticles were synthesized by nanoprecipitation together with the near-infrared dye indocyanine green (ICG) for particle stabilization and in vivo tracing (fig. S1D). Control nanoparticles were produced using dextran (Dex), which is biochemically similar to fucoidan but does not bind P-selectin (DexNav and DexdBET6) (*48*). All SMNPs and Dex nanoparticle formulations exhibited uniform size distributions, similar hydrodynamic diameters, similar surface charge properties, and stability (fig. S1E-G).

To assess biodistribution in vivo, SMNPs or control nanoparticles were administered intravenously to mice with established liver or lung fibrosis, and ICG signal was quantified in 6 vital organs at 12 or 24 hours post-treatment. In the CCl_4_-induced liver fibrosis model, FiNav and FidBET6 nanoparticles preferentially accumulated in the liver, whereas in the bleomycin-induced lung fibrosis model, SMNPs selectively enriched in the lung (Fig. 1I and fig. S1H, I). By contrast, control DexNav and DexdBET6, which do not bind P-selectin, showed no organ-specific accumulation (Fig. 1I and fig. S1H, I). At the cellular level, FiNav and FidBET6 nanoparticles were enriched within P-selectin^+^/SA-β-gal^+^ cells (Fig. 1J, K and fig. S1J). Together, these data establish that P-selectin can be leveraged to selectively deliver senescence-modulating payloads to senescent cells in fibrotic organs, providing a foundation for evaluating therapeutic efficacy and safety.

### SMNPs reverse established fibrosis while limiting systemic toxicities

Prior work has shown that Navitoclax and BET inhibitors can attenuate fibrosis in preclinical models (*15, 61, 62*), but their clinical development in fibrosis and cancer has been impeded by dose-limiting hematologic and systemic toxicities (*38, 39, 63–67*). We therefore asked whether P-selectin-directed SMNPs could preserve the antifibrotic efficacy of these agents while mitigating toxicity. To this end, we performed treatment studies in CCl_4_- and bleomycin-induced models of liver and lung fibrosis, respectively (Fig. 2A-B). After fibrosis was established, mice were treated with vehicle, SMNPs (FiNav or FidBET6), control nanoparticles (DexNav or DexdBET6), matched doses of free Navitoclax or dBET6, or free fucoidan. Treatment efficacy was evaluated by quantifying senescent cells, collagen deposition (Sirius red), and α-SMA^+^ activated fibroblasts, with portal vein flow (*68*) and serum enzymes (ALT/AST) as additional measures of liver function. Systemic toxicities were assessed by hematologic profiling, body-weight monitoring, and treatment-associated mortality.

**Fig. 2:**
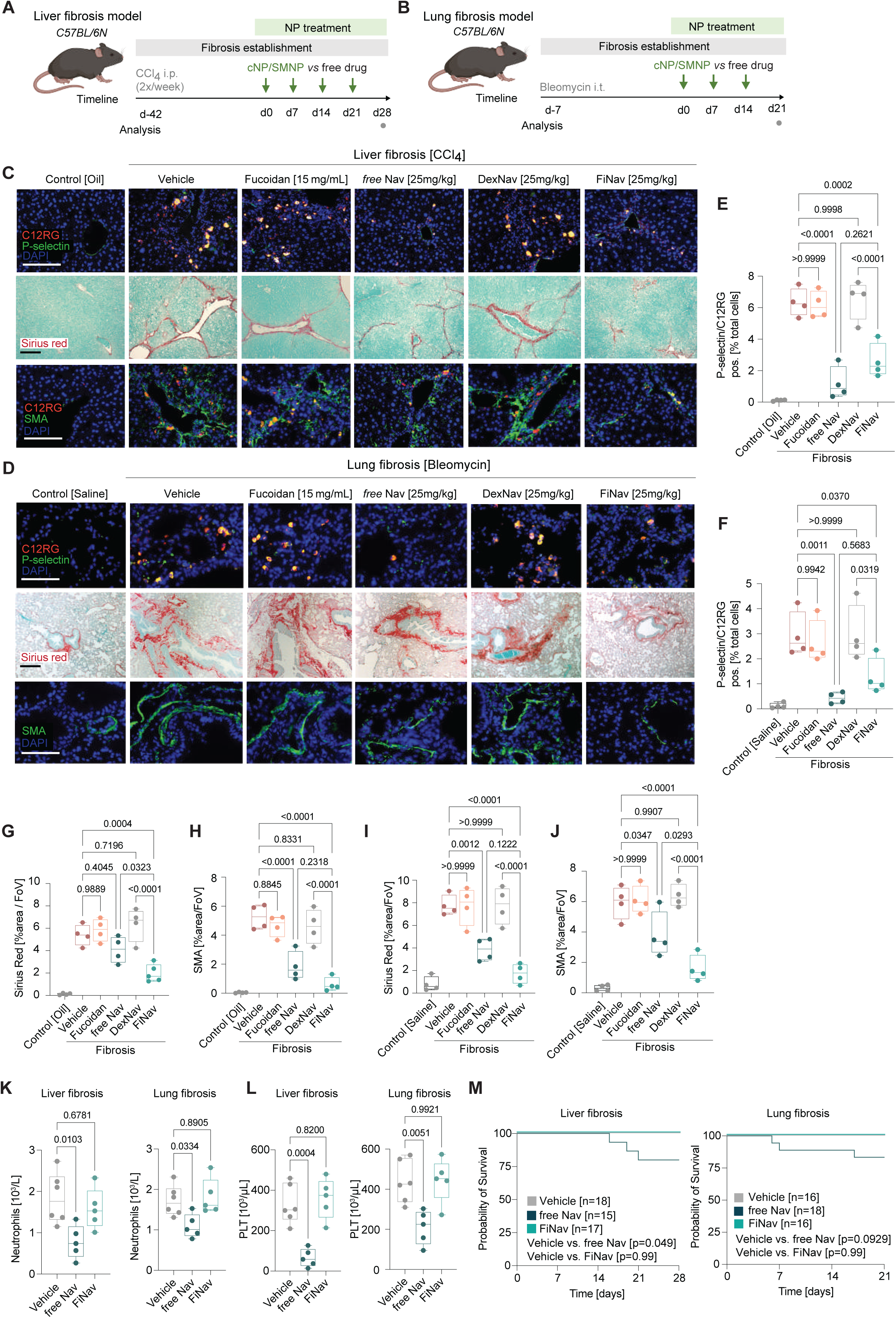
FiNav nanoparticles selectively deplete senescent cells and attenuate fibrosis with minimal toxicity. (**A,B**) Schematic of nanoparticle (NP) treatment schedules in murine fibrotic livers (A) and lungs (B); cNP: control nanoparticles. (**C-J**) Representative images of P-selectin^+^ senescent (SA-β-gal^+^ per C12RG assay) cells, collagen deposition (Sirius Red), and myofibroblast activation (SMA) in fibrotic livers (C) and lungs (D) treated as indicated, with corresponding quantifications (livers: E, G, H and lungs: F, I, J) (4-6 mice/group). Scale bars, 150 μm. (**K-L**) Neutrophil (K) and platelet (L) counts from blood of mice with fibrotic livers (left) or lungs (right) treated with free navitoclax or FiNav nanoparticles (5-6 mice/group). (**M**) Toxicity of free navitoclax and FiNav assessed by Kaplan-Meier survival curves for mice with fibrotic livers (left) or fibrotic lungs (right) treated with free navitoclax or FiNav nanoparticles. Each dot represents one mouse. Data are presented as mean ± s.e.m. Statistical tests were one-way ANOVA with Sidak’s multiple comparisons test (E-L) and log-rank Mantel Cox test (M).

Across both liver and lung fibrosis models, FiNav treatment efficiently reduced P-selectin^+^/SA-β-gal^+^ double-positive cells, whereas FidBET6 had little or no effect on this compartment (Fig. 2C-F and fig. S2A-D). Despite these differences, both FiNav and FidBET6 significantly reduced fibrotic burden, as evidenced by decreased collagen deposition and reduced numbers of α-SMA^+^ activated fibroblasts (Fig. 2C-D, G-J and fig. S2A-B, E-H). This effect was at least as great as that achieved with free drug, despite SMNPs targeting only a small fraction of cells in tissue (Fig. 2G–H and fig. S2E–H). Control nanoparticles harboring either Nav or dBET6 were ineffective, establishing a requirement for P-selectin–dependent targeting for nanoparticle-based therapeutic benefit. Importantly, at similar doses used for SMNP production, free fucoidan had no effect.

SMNPs also improved organ function. In mice with liver fibrosis, SMNPs improved portal vein flow, with FiNav producing a significant advantage over free Navitoclax, whereas free fucoidan and control nanoparticles had no measurable impact (fig. S2I). Additionally, SMNPs significantly reduced liver damage, as assessed by serum ALT and AST levels (fig. S2J,K). These findings indicate that nanoencapsulation preserves antifibrotic efficacy and may confer additional functional benefits in restoring organ physiology.

Finally, we evaluated whether SMNP-mediated delivery mitigates the dose-limiting toxicities associated with Navitoclax and BRD4-targeting agents, including neutropenia and thrombocytopenia. In both liver and lung fibrosis models, FiNav and FidBET6 treatment eliminated these toxicities (Fig. 2K,L and fig. S3A,B). Consistent with preserved platelet function, SMNP-treated mice exhibited significantly shorter bleeding times compared to free-drug–treated animals (fig. S3C,D). Extended toxicity profiling further revealed that SMNPs produced with either Navitoclax or dBET6 protected from free drug-associated leukopenia, lymphopenia, body-weight loss, and treatment-associated mortality (Fig. 2M and fig. S3E-M). Therefore, SMNPs harboring either a small molecule senolytic or senomorphic drug can reverse established fibrosis in both liver and lung while markedly improving the therapeutic index.

### SMNPs target a subset of senescent-like macrophages

To better define the cell types and states targeted by SMNPs in vivo, we performed single-cell RNA sequencing (scRNA-seq) on cells isolated from murine fibrotic liver and lung tissues. Owing to variability in cellular autofluorescence across cell types and the low fraction of SMNP target cells, fluorescence activated cell sorting (FACS) of cells based on their SMNP-associated ICG signal was not consistently achievable. Based on our observation that SMNPs preferentially accumulate in P-selectin/SA-β-gal double-positive cells (Fig. 1J,K and fig. S1J), we instead leveraged endogenous P-selectin surface expression in combination with a fluorescent SA-β-gal probe (SPiDER, functionally analogous to C12FDG (*55, 69*) to enrich for potential target populations (fig. S4A). Following organ digestion, four distinct cell populations were isolated by FACS: P-selectin^+^/SPiDER^+^ (P⁺sen⁺), P-selectin^+^/SPiDER⁻ (P⁺sen⁻), P-selectin⁻/SPiDER^+^ (P⁻sen⁺), and P-selectin⁻/SPiDER⁻ (P⁻sen⁻). scRNA-seq analysis revealed that the P⁺sen⁺ compartment from both liver and lung was composed predominantly of macrophages (*Adgre1⁺, Cd68⁺, Csf1r⁺*) and endothelial cells (*Pecam1⁺, Cdh5⁺, Icam1⁺*), with smaller contributions from fibroblasts and epithelial cells in the fibrotic lung (Fig. 3A–D; fig. S4B–G). The identities of P⁺sen⁺ macrophages and endothelial cells inferred by scRNA-seq were validated in situ using C12RG-based SA-β-gal staining combined with immunofluorescence for macrophage (CD68, F4/80) and endothelial (CD31) markers (Fig. 3E,F; fig. S4H).

**Fig. 3:**
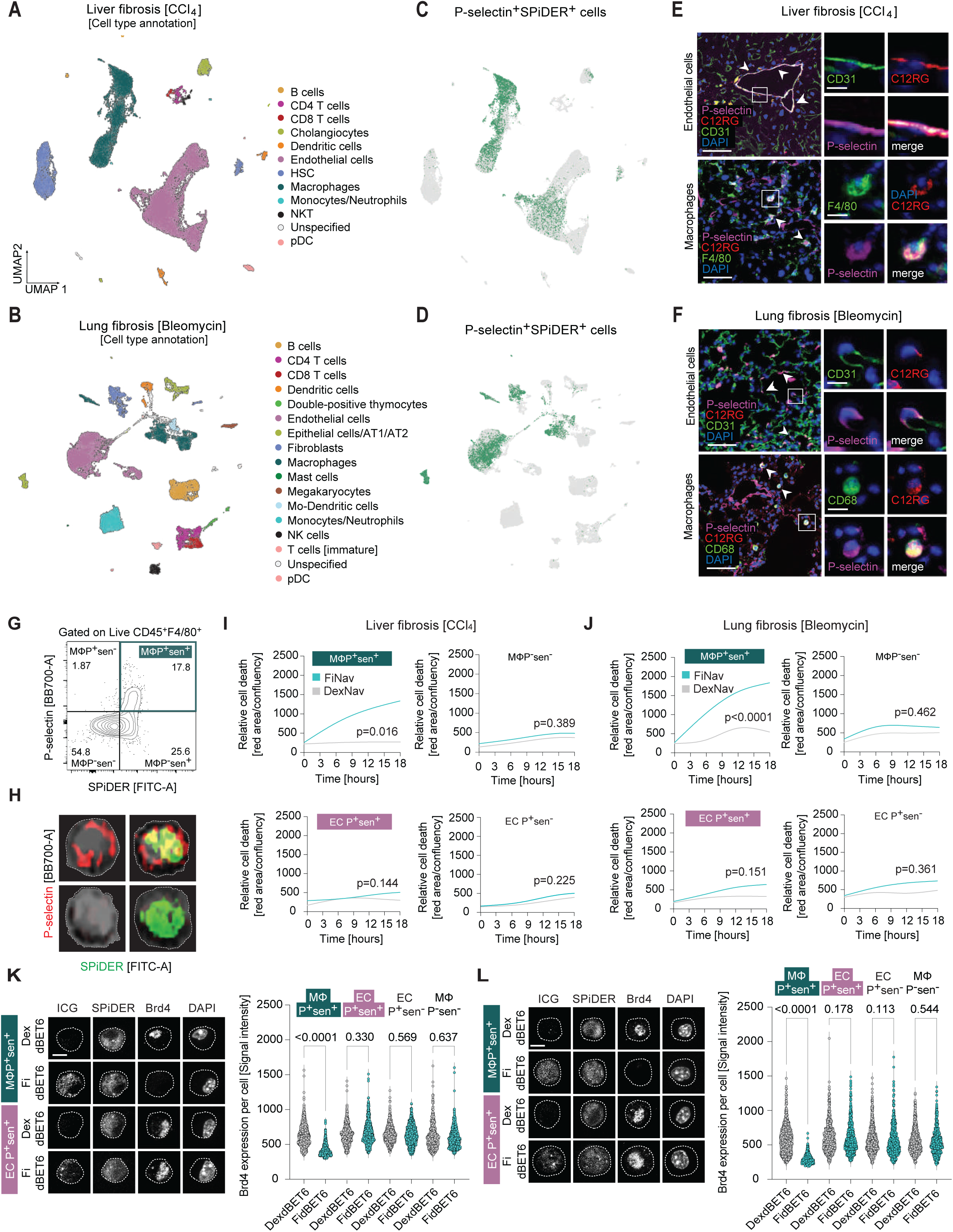
SMNPs target a P-selectin/SA-β-gal double-positive macrophage subpopulation in fibrotic liver and lungs. (**A-B**) Visualization of unsupervised clustering in a UMAP of cells isolated from murine fibrotic (A) livers and (B) lungs. **(C-D)** UMAP visualization of P^+^sen^+^ cells (green) in A and B. (**E-F**) Representative IF images of SA-β-gal (C12RG^+^) and P-selectin expression in CD31^+^ endothelial cells (EC), F4/80^+^ (liver) and CD68^+^ (lung) macrophages of murine fibrotic (E) livers and (F) lungs. Scale bars, 100 μm. (**G**) Representative gating on Live/Dead^-^ CD45^+^ F4/80^+^ cells based on P-selectin expression and SA-β-gal activity (SPiDER). (**H**) Surface P-selectin expression (red) and SPiDER activity (green) in MΦ populations sorted in (G), visualized by BD FACSDiscover S8 Cell Sorter. (**I-J)** Ex-vivo killing of sorted P^+^sen^+^or P^+^sen^-^ ECs and P^+^sen^+^ or P^-^sen^-^ MΦs from murine fibrotic (I) livers and (J) lungs treated with DexNav or FiNav. Representative data from three independent experiments. **(K-L)** IF images (left panels) of nanoparticles (ICG), SPiDER activity, and Brd4 in sorted P^+^sen^+^ EC and MΦ from murine fibrotic livers (K) or lungs (L) following treatment with DexdBET6 or FidBET6 (left), and quantification of Brd4 (right) (three biological replicates). Scale bars, 5 μm. Each dot represents one cell. Data are presented as mean ± s.e.m. Mann Whitney test or Welch’s t test were used at endpoint (18h) for (I,J), and Mann Whitney test for (K,L).

To directly establish the SMNP-targeted populations, we assessed SMNP uptake and pharmacologic activity across senescent-like and non-senescent subsets of macrophage and endothelial lineages. Using a modified FACS strategy incorporating P-selectin surface expression, SPiDER, and lineage-specific markers, we isolated P⁺sen⁺, P⁺sen⁻, P⁻sen⁺, and P⁻sen⁻ subsets within macrophage and endothelial compartments from fibrotic liver and lung (Fig. 3G,H, fig. S5A). Macrophages were defined as CD45^+^/F4/80^+^ cells in the liver and CD45^+^/CD64^+^ cells in the lung, whereas endothelial cells were defined as CD45⁻/CD31^+^ (fig S5A). Purified populations (fig. S5A,B) were subjected to short-term ex vivo treatment with Navitoclax-based or dBET6-based nanoparticles. Targeting was evaluated using Incucyte-based cell viability assays for cell preparations treated with Navitoclax-based nanoparticles or Brd4 immunofluorescence for those treated with dBET6-based nanoparticles.

These studies pinpointed a P⁺sen⁺ macrophage population (MΦP^+^sen^+^) as the principal cellular target of SMNPs in both liver and lung fibrosis. Accordingly, FiNav selectively induced cell death in MΦP^+^sen^+^ while sparing all other macrophage and endothelial subsets irrespective of P-selectin or SPiDER status (Fig. 3I, J and fig. S5 C,D). Consistent with this selectivity, complementary analysis with FidBET6 revealed selective degradation of Brd4 protein within MΦP^+^sen^+^ cells (Fig. 3K, L and fig. S5E, F). By contrast, neither P^+^sen^+^ nor P^-^sen^-^ endothelial populations (ECP^+^sen^+^ and ECP^-^sen^-^ respectively) showed evidence of functional perturbation by either SMNP. Extending these observations in vivo, FiNav treatment led to depletion of MΦP^+^sen^+^ populations in liver and lung within 36 hours, with minimal effects on the other populations analyzed (fig. S5G–I). Notably, depletion of MΦP^+^sen^+^ populations preceded fibrosis resolution, supporting the idea that MΦP^+^sen^+^ depletion is a cause rather than a consequence of tissue remodeling (fig. S5J).

We next asked why SMNPs selectively impact MΦP^+^sen⁺ cells. To test whether P-selectin mediates uptake, we added a P-selectin–blocking antibody to sorted MΦP^+^sen⁺ and ECP^+^sen⁺ cells. This markedly reduced the SMNP-associated ICG signal, supporting P-selectin-dependent targeting rather than nonspecific uptake (fig. S6A). However, P-selectin expression alone did not explain functional selectivity. Although initial SMNP uptake was comparable in MΦP^+^sen⁺ and ECP^+^sen⁺ cells, SMNP-associated ICG fluorescence was retained after washout only in MΦP^+^sen^+^ cells (fig. S6B). Subcellular localization studies showed that endothelial cells routed SMNPs to Caveolin-1^+^ vesicles and Rab5^+^ early endosomes, consistent with prior studies showing caveolin-dependent transcytosis of fucoidan-based nanoparticles (*46–48*). However, MΦP^+^sen⁺ macrophages routed SMNPs to Rab7^+^ late endosomes and Lamp1^+^ lysosomes, a traffic pattern consistent with efficient intracellular payload release (fig. S6C,D). In agreement, lysosomal gene expression programs were enriched in macrophages relative to endothelial cells isolated from fibrotic liver and lung (fig. S6E,F).

To determine whether SMNP selectivity also requires a senescence-targeting payload, we generated dextran- and fucoidan-based nanoparticles encapsulating Paclitaxel (DexPax and FiPax), a broadly cytotoxic agent not linked to senescence-associated pathways. Paclitaxel-loaded particles had similar size and charge to Navitoclax particles (fig S7A-C) and showed the expected P-selectin-mediated uptake following treatment of macrophages ex vivo (fig. S7D). As expected, free Navitoclax was more toxic in MΦP^+^sen⁺ and MΦP^−^sen⁺ cells than in MΦP⁺sen^-^ and MΦP^-^sen^-^ cells, whereas free Paclitaxel was active across all macrophage subsets (Fig. S7E–H). Consistently, FiPax killed P-selectin^+^ macrophages irrespective of senescence state, whereas FiNav remained selective for MΦP⁺sen⁺ cells (fig S7E-H). In vivo, however, FiPax showed reduced anti-fibrotic activity and did not protect from Paclitaxel toxicity (fig. S7I–Q). Thus, SMNP selectivity reflects the convergence of P-selectin–dependent delivery, cell-type-specific intracellular routing, and payload activity matched to senescence-state vulnerabilities.

### SMNP-targeted macrophage populations are related to lipid-associated macrophages

The identification of a senescent-like macrophage population as a key driver of fibrosis is consistent with studies demonstrating that genetic ablation of p16^+^ macrophages reverses liver fibrosis in mice (*36*). Additional work implicates pathogenic roles for p16^+^ macrophages in lung disease (*29, 70*), collectively supporting a conserved role for such macrophages in tissue pathology across organs. To define the molecular identity and functional properties of this population, we performed differential gene expression analysis using Model-based Analysis of Single-cell Transcriptomics (MAST) (*71*) by comparing MΦP^+^sen^+^ to MΦP⁻sen⁻ macrophages to generate a MΦP^+^sen^+^ signature.

In the liver, MΦP^+^sen^+^ macrophages displayed a robust senescence-associated transcriptional program, including upregulation of *Cdkn1a, Cdkn2a,* and *Jun*, induction of SASP-associated factors (*Cxcl1, Cxcl2, Il1a, Mmp27*), and downregulation of *Lmnb1* and *Ccnd2* (fig. S8A). Pathway analysis revealed enrichment for p53 signaling and lysosomal function and depletion of cell-cycle progression, E2F targets, and DNA repair pathways (fig. S8B) (*72–74*). MΦP^+^sen^+^ macrophages isolated from fibrotic lungs showed a concordant transcriptional profile, including upregulation of *Cdkn1a, Il1a, Cxcl2*, *Mmp12*, downregulation of *Lmnb1, E2f2*, *Ccnd3*, enrichment of p53 and lysosomal pathways, and suppression of programs linked to apoptosis, cell-cycle regulation, and DNA repair (fig. S8C, D).

MΦP^+^sen^+^ cells were strongly enriched for the consensus senNET senescence signature (*49*) as well as transcriptional signatures derived from p16^+^ pathogenic macrophages in fibrotic livers (*36*) and lung tumorigenesis (*29, 70*) (Fig. 4A-H and fig. S8E-H). Nevertheless, this analysis revealed that senescent programs are not restricted to MΦP^+^sen^+^ but are also enriched in other macrophage subtypes. Indeed, mIF identified macrophage subsets that expressed senescence markers without P-selectin. However, most P-selectin^+^ macrophages co-expressed senescence markers (fig. S8I,J).

**Fig. 4:**
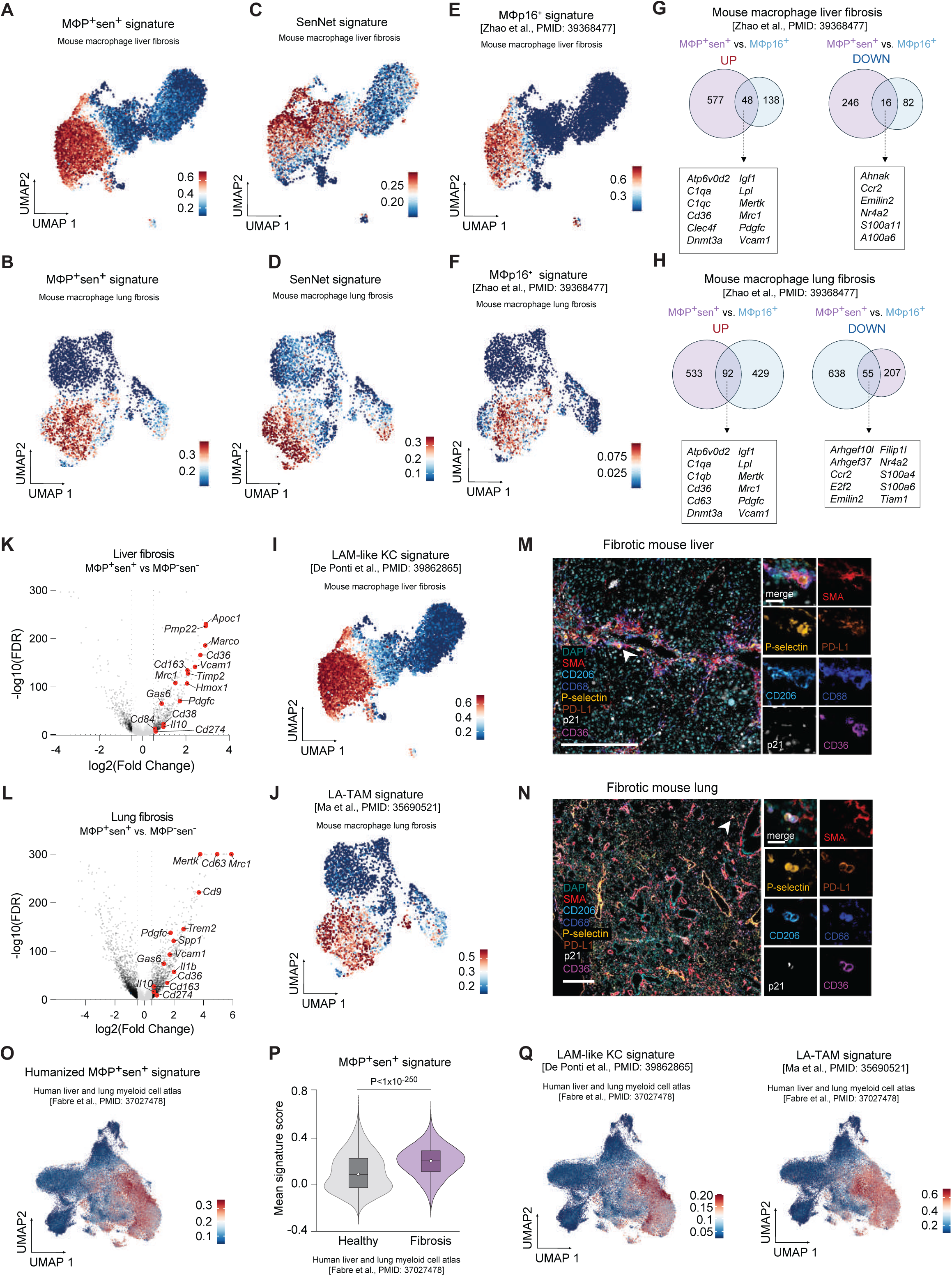
P-selectin/SA–β-gal double-positive macrophages exhibit senescence and LAM-related programs and shape immunosuppressive niches. (**A**-**F**) UMAP visualization of gene signature scores. Cells are scored for MΦP^+^sen^+^ (A-B), SenNet senescence (C-D), and published p16^+^ macrophage (E-F) signatures in murine fibrotic liver (A, C, E) and lung (B, D, F). (**G-H**) Venn diagrams representing the common upregulated (left) or downregulated (right) genes in MΦP^+^sen^+^ compared to those in p16^+^ macrophages in murine liver (G) and lung (H) fibrosis. (**I-J**) Volcano plot showing differentially expressed genes in MΦP^+^sen^+^ compared to MΦP^-^sen^-^ from murine liver (I) and lung (J) fibrosis, with selected pro-fibrotic and immunosuppressive genes labeled. (**K-L**) UMAP visualization of gene signature scores. Cells are scored for the lipid-associated macrophage signature in murine liver (K) and lung (L) fibrosis. (**M-N**) Representative mIF images showing co-expression of immunosuppressive markers (CD206, PD-L1) in P-selectin^+^ macrophages (CD68^+^) expressing a senescence marker (p21^+^) in murine liver (M) and lung (N) fibrosis. Scale bars, 200 μm and 20 μm (magnification). (**O**) Humanized MΦP^+^sen^+^ macrophage signature in pooled scRNA datasets of human liver and lung fibrosis samples. (**P**) Violin plot of humanized MΦP^+^sen^+^ signature score in human healthy and fibrotic liver and lungs. (**Q**) UMAP visualization of LAM signatures in human liver and lung fibrosis dataset. Wilcoxon rank-sum test was used for (P).

In addition to senescence-associated features, MΦP^+^sen^+^ macrophages displayed hallmarks of lipid-associated macrophages (LAMs), which have been linked to lipid handling, matrix remodeling, chronic inflammation, and immune suppression (*75–78*). Across both organs, MΦP^+^sen^+^ cells were enriched for lipid, fatty acid and cholesterol metabolism gene signatures, and depleted for antigen processing and presentation programs (fig. S8B,D). MΦP^+^sen^+^ cells upregulate a plethora of immunomodulatory and immune-suppressive genes (*Cd274*, *Mrc1*, *Gas6*, *Hmox1*, *Spp1*, *Cd38*, and *Il10*) and genes involved in matrix remodeling (*Cd36*, *Pdgfc*, *Il1a*, *Marco*, *Timp2*, *Cd5l*, *Cd163*, *Vcam1*), many of which are hallmark features of LAMs (*75–77*) (Fig. 4I,J). Consistently, MΦP⁺sen⁺ transcriptional programs showed strong overlap with published LAM signatures (Fig. 4K,L) (*76, 78*). Using mIF, we orthogonally validated LAM-associated proteins including CD36 and CD206 (encoded by *Mrc1*) and immune-suppressive marker PD-L1 (encoded by *Cd274*) on MΦP^+^sen^+^ cells (defined by P-selectin^+^, p21^+^, macrophage markers CD68^+^ and/or F480^+^, and LaminB1^-^ and Ki67^-^) (Fig. 4 M,N). However, as with senescence-associated transcriptional signature, LAM-associated programs were not exclusive to MΦP⁺sen⁺ cells. mIF analysis demonstrated that CD36, a canonical LAM-associated scavenger receptor on macrophages, accumulates in fibrosis, with only a subset of these cells co-expressing P-selectin (fig. S8K).

To assess conservation in human disease, we analyzed publicly available scRNA-seq datasets from human liver (44 healthy and 5 cirrhotic patients across five studies) and lung (51 healthy and 66 fibrotic patients across four studies) (*79*). This analysis identified a macrophage population that was expanded in fibrosis and enriched for both the humanized MΦP^+^sen^+^ and LAM-associated gene expression signature (Fig. 4O-Q). Thus, MΦP^+^sen^+^ macrophages define a fibrosis-enriched, senescent-like macrophage state with LAM-like features that is conserved across murine models and human fibrosis.

### SMNPs remodel the immune landscape of fibrotic tissues

To define the spatial organization MΦP^+^sen^+^ populations within fibrotic tissues, we examined their local cellular neighborhoods in situ. mIF of fibrotic liver and lung, coupled with spatial neighborhood analysis using MΦP⁺sen⁺ cells as anchor points, revealed preferential localization adjacent to activated α-SMA⁺ fibroblasts and endothelial cells in both organs, defining a conserved pro-fibrotic stromal niche. In contrast, antigen-presenting and adaptive immune subsets, including dendritic cells (DCs), conventional CD4⁺ and CD8⁺ T cells, and B cells, were largely excluded from these regions, consistent with localized immune suppression (fig. S8L,M).

To determine how SMNP-mediated targeting impacts these niches, fibrotic livers and lungs were isolated from mice following treatment with SMNPs (FiNav or FidBET6) or control nanoparticles (DexNav or DexdBET6) and were analyzed by mIF for immune cell markers. SMNP treatment induced profound tissue remodeling associated with an overall reduction in macrophage content, shifting the residual macrophage composition from anti-inflammatory phenotypes (MHCII⁻CD206⁺ and MHCII⁻LYVE1⁺PD-L1⁺) toward proinflammatory/antigen-presenting cell (APC) states (MHCII⁺CD11c⁺ and MHCII⁺CD206⁺) (fig. S9A-H). Concomitantly, SMNP treatment expanded CD4⁺ and CD8⁺ T and B cell populations within regions once dominated by profibrotic stroma (fig. S9A-H). Neither free drugs nor control nanoparticles produced significant recruitment of CD3+ T cell populations, indicating that targeting a P-selectin population is required (fig. S10A,B). In parallel, SMNP treatment promoted an influx of DCs (CD45^+^MHCII⁺CD11c⁺ and CD68^-^ and F4/80^-^), accompanied by a shift from tolerogenic (tolDC; [MHC-II^low^ ^or-^/PD-L1^+^]), a DC subtype involved in immune tolerance, towards a maturated/activated (CD80^+^/CD86^+^/CD40^+/-^) DC phenotype (fig. S10C,D).

Consistent with predicted depletion of MΦP⁺sen⁺ cells, cytokine profiling of fibrotic liver and lung tissues after SMNP treatment show reduction of pro-fibrotic and immunosuppressive mediators (including amphiregulin, CCL5, IL-1β) while inducing cytokines associated with increased immune response (including IL-2, IL-4, IL-7, and IFN-γ) (fig. S10E–H). Together, these coordinated cellular and secretory changes indicate that SMNPs resolve fibrosis while reprogramming immunosuppressive niches toward immune-permissive microenvironments.

### SMNPs can reverse immunotherapy resistance

Many solid tumors which arise within, or actively induce, a fibrotic stroma develop immune-excluded microenvironments that limit T cell infiltration and confer resistance to immune checkpoint blockade (ICB) (*8–11*). This is particularly evident in liver and lung cancers, which frequently arise within chronically inflamed, fibrotic organs, where stromal desmoplasia and myeloid-driven immune suppression are strongly linked to poor clinical outcomes (*10, 11*). Although immune-suppressive macrophage populations are central mediators of these barriers, strategies to selectively target them and reprogram the fibrotic TME remain limited (*76, 80*). Based on our findings that MΦP⁺sen⁺ macrophages sustain fibrotic, immune-excluded niches and are efficiently engaged by SMNPs, we tested whether SMNP-mediated remodeling of the fibrotic TME could restore ICB sensitivity.

To test this, we engineered murine models of liver and lung cancer that respond to clinically relevant first line ICB-containing therapies when engrafted into normal organs and engrafted these into mice in which liver or lung fibrosis was established by CCl_4_ or bleomycin, respectively (fig. S11A-B). Liver tumors were engrafted by orthotopic transplantation of syngeneic *Myc-GFP; Trp53^-/-^* (MP) hepatocellular carcinoma (HCC) tumor cells (*81*) (Fig. 5A), whereas lung tumors were established by intravenous injection of Kras^G12D/+;^ Trp53^-/-^ (KP) non-small cell lung cancer (NSCLC) tumor cells (*82*) (Fig. 5B). Although our tumor models did not express P-selectin in vitro, to avoid any potential consequences of tumor-upregulation of P-selectin expression in vivo, we took the added precaution of silencing P-selectin using validated shRNAs to avoid potential confounding effects on data interpretation (fig. S11C). Tumor-bearing mice were randomized by baseline imaging and treated with SMNPs to remodel the fibrotic niche before and during administration of clinically relevant, first-line ICB-based regimens: anti-PD-L1 combined with anti-VEGF in liver tumors, or anti-PD-L1 combined with cisplatin in lung tumors (hereby referred to as Tx). Tumor growth, survival, and immune responses were monitored longitudinally (Fig. 5A, B).

**Fig. 5:**
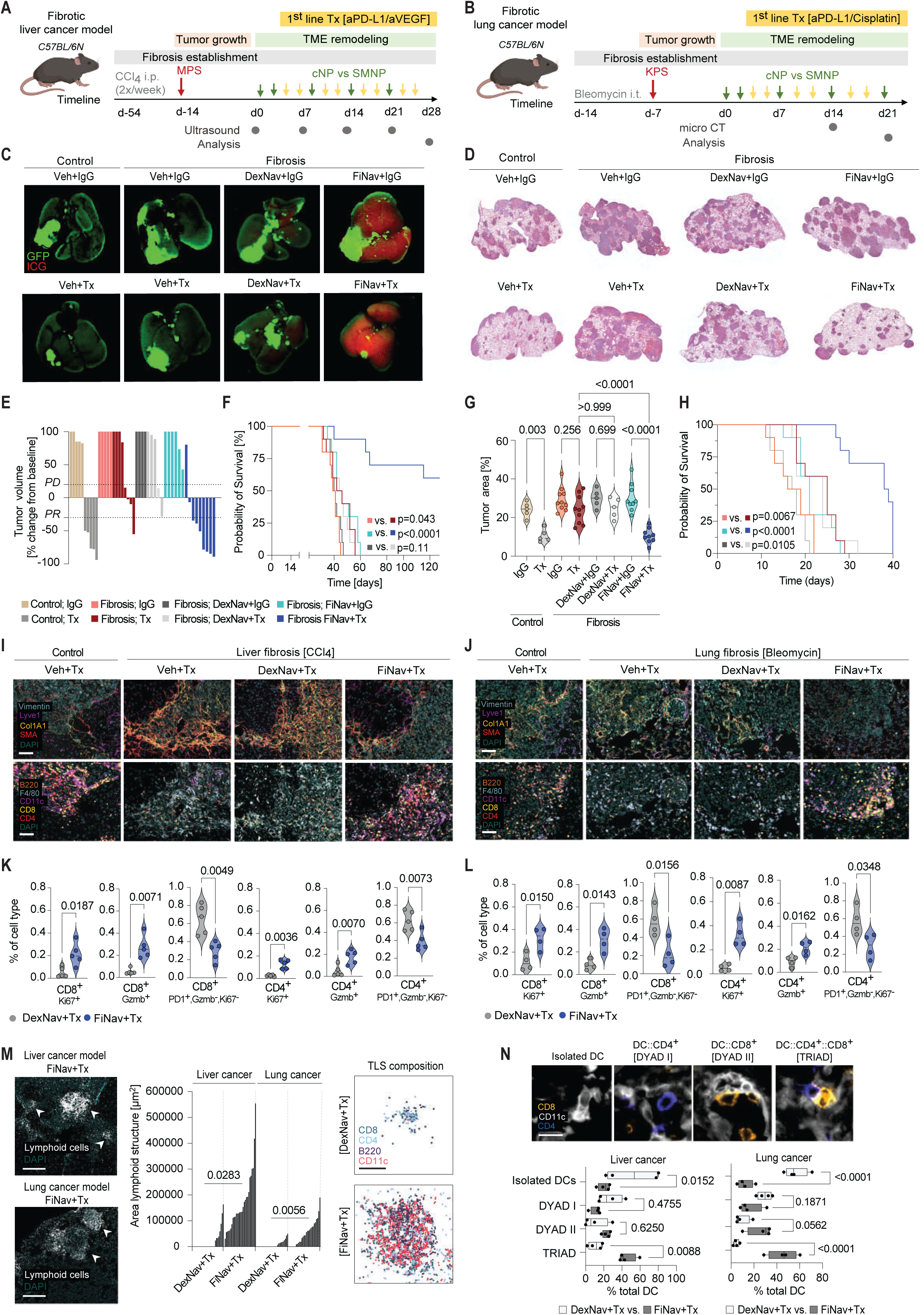
SMNP-mediated niche remodeling overcomes resistance to immune checkpoint blockade. (**A**-**B**) Treatment schedule for murine fibrotic liver (A) and lung (B) tumor models treated with nanoparticles (cNP; SMNP) and first-line therapy (Tx). MPS: *Myc-GFP;p53^KO^;shSelp* murine liver cancer cells. KPS: *Kras^G12D^;p53^KO^; shSelp* murine lung cancer cells. (**C-D**) Representative GFP/ICG fluorescence imaging of liver tumors (C) and H&E macroscopic sections of lung tumors (D) after DexNav or FiNav plus Tx. (**E-F**) Tumor volume change between d0 and d14 (5-10 mice/group) (E) and overall survival (F) (10 mice/group) of liver tumors model with or without fibrosis induction treated with DexNav or FiNav plus Tx. (**G-H**) Tumor area at d21 (5-10 mice/group) (G) and overall survival (10 mice/group) (H) of mice with or without fibrosis induction with lung tumors treated with DexNav or FiNav plus Tx. (**I**-**J**) mIF analysis of stromal (Vimentin, Lyve1, Col1a1, SMA) and immune (B220, F4/80, Cd11c, CD8, CD4) markers in murine fibrotic liver (I) or lung (J) tumors treated with DexNav or FiNav plus Tx. Scale bars, 100 μm. (**K**-**L**) Quantification of active/cytotoxic/exhausted CD8^+^ and CD4^+^ T cells in murine fibrotic liver (K) or lung (L) tumors treated with DexNav or FiNav plus Tx (5 mice/group). (**M**) (Left) Representative IF images of lymphoid aggregates (CD3, CD8, CD4, B220 markers in white) in fibrotic liver (top) and lung (bottom) cancer models treated with FiNav plus Tx and (middle) quantification (4 mice/group). (Right) Spatial plot showing CD8^+^ T-cells, CD4^+^ T-cells, B220^+^ B-cells, and CD11c^+^ dendritic cells of murine fibrotic liver and lung cancer treated with DexNav or FiNav plus Tx. (**N**) (Top) Representative IF pictures of isolated DC (CD11c^+^), DC::CD4 dyads (DYAD I) (CD11c^+^::CD4^+^), DC::CD8 dyads (DYAD II) (CD11c^+^::CD8^+^) and DC::CD4:CD8 triads (TRIAD) (CD11c^+^::CD4^+^::CD8^+^) in murine fibrotic liver and lung cancer treated with DexNav or FiNav plus Tx. (Bottom) Quantification of isolated DCs, DYADIs, DYADIIs, and TRIADs in murine fibrotic liver (left) and lung (right) cancer treated with DexNav or FiNav plus Tx (4 mice/group). Individual bars (E) or dots (G, K-L, N) each represent one mouse. Individual bars (M) represent one lymphoid aggregate area quantified from 4 mice/group. Data are presented as mean ± s.e.m. Statistical tests were log-rank Mantel Cox test (F, H), one-way ANOVA with Sidak’s multiple comparisons (G, N), and Mann Whitney or Welch’s t test (K, L, M).

Mirroring resistance patterns observed in patients (*10, 83*), MPS and KPS tumors that responded robustly to Tx in non-fibrotic liver and lung were refractory when embedded within fibrotic TMEs (Fig. 5C-E, G). SMNP monotherapy with FiNav induced modest tumor growth inhibition and survival benefits, whereas control nanoparticles (DexNav) were ineffective (Fig. 5C-H). Strikingly, combining FiNav+Tx produced durable tumor regression in both fibrotic tumor models, accompanied by significant improvements in overall survival (Fig. 5C-H and fig. S11 D, E). Comparable therapeutic synergy was observed using FidBET6 in the liver cancer model (fig. S11F), demonstrating that both senolytic and senomorphic SMNPs can sensitize fibrotic tumors to immunotherapy.

To define how SMNPs remodel the fibrotic TME, we performed mIF profiling of MPS and KPS tumors and adjacent liver and lung stroma, respectively (Fig. 5I, J). Untreated fibrotic tumors were characterized by expanded αSMA^+^ activated fibroblasts, dense collagen I deposition, increased LYVE1^+^ lymphatic endothelium, and CD68^+^ or F4/80^+^ macrophage accumulation at the tumor border, with concomitant exclusion of DCs and CD4^+^ and CD8^+^ T cells (Fig. 5 I,J and fig. S12 A-H). This architecture persisted in DexNav-treated mice (Fig. 5I, J and fig. S12 A-D). In contrast, FiNav alone dismantled these stromal barriers, reducing collagen type I deposition, αSMA^+^ myofibroblasts, and macrophage density. Concordantly, spatial neighborhood analysis at the tumor border revealed the collapse of the pro-fibrotic, immune-suppressive niche and its replacement by DCs, B cells, and effector T cells (fig. S12 I,J).

Despite extensive remodeling of the fibrotic TME, FiNav monotherapy was insufficient to achieve durable tumor control, indicating that niche disruption alone does not fully restore anti-tumor immunity. Accordingly, immunophenotyping revealed that, while FiNav produced a modest increase in activated and proliferating CD4^+^ and CD8^+^ T cells, the majority of T cell populations displayed markers of exhaustion (fig. S12K,L). By contrast, combining FiNav with ICB based therapies triggered a marked expansion of cytotoxic and proliferating T cells, increased conventional CD4^+^ T cells, and concomitantly reduced exhausted T cell and Treg populations in both liver and lung tumors (Fig. 5K,L, fig S12K,L, fig S13A). These results establish that SMNP-mediated niche reprogramming creates a permissive context in which ICB can elicit robust and sustained anti-tumor immunity.

Consistent with the restoration of effector immune programs, FiNav+Tx induced spatially organized lymphoid aggregates containing DCs, B cells, and CD4^+^ and CD8^+^ T cells in fibrotic liver and lung tumors (Fig. 5M, fig. S13B-C). DC-focused mIF further showed fewer isolated DCs and increased coordinated DC cell interactions, including DC:CD4:CD8 immune triads (Fig. 5N, fig. S13D), which are associated with productive antigen presentation and local T cell effector function (*84*). Concordantly, FiNav+Tx treatment reduced tolDCs and shifted DC composition toward mature and activated states (fig. S13E-F). Consistent with the potential of such structures to promote anti-tumor immunity, antibody-mediated depletion of CD8^+^ T cells largely abrogated the survival benefit of FiNav+Tx, CD4^+^ T cell depletion produced a more modest loss of benefit, and NK cell depletion had minimal impact (fig. S13G-I). Thus, SMNP-mediated niche remodeling creates organized, antigen-presenting immune microenvironments that support CD8^+^ and CD4^+^ T-cell-dependent tumor control.

### SMNPs target a conserved senescent-like macrophage state in human tumors

To assess whether macrophages analogous to the MΦP^+^sen^+^ populations identified in fibrotic murine tissues are present in human liver and lung tumors, we applied the humanized MΦP⁺sen⁺ gene signature to published scRNA-seq datasets from treatment-naïve HCC (n=79) (*85*) and NSCLC (n=12) (*86*). In both tumor types, the MΦP⁺sen⁺ signature identified a distinct myeloid subcluster (Fig. 6A, B). Mapping this signature onto a previously described framework stratifying the HCC tumor immune microenvironment (TIME) into distinct stages ranging from immune activation to immune suppression revealed preferential alignment with the myeloid-driven immune-excluded niche (fig. S14A) (*85*). Comparison of the MΦP⁺sen⁺ signature with established tumor-associated macrophage programs (*78*) revealed strongest overlap with Kupffer-like TAMs, interstitial resident tissue macrophages, and LAMs (Fig. 6C, D), populations previously implicated in immune suppression (*78, 87–89*). Consistent with our murine data demonstrating that MΦP^+^sen^+^ cells represent a distinct macrophage subset within the broader LAM compartment, CD36 expressing macrophages were present in both the low-fibrosis and high-fibrosis human lung cancer TMEs, but P-selectin^+^/CD36^+^ subsets preferentially accumulated in the high-fibrosis setting (fig. S14B-D).

**Fig. 6:**
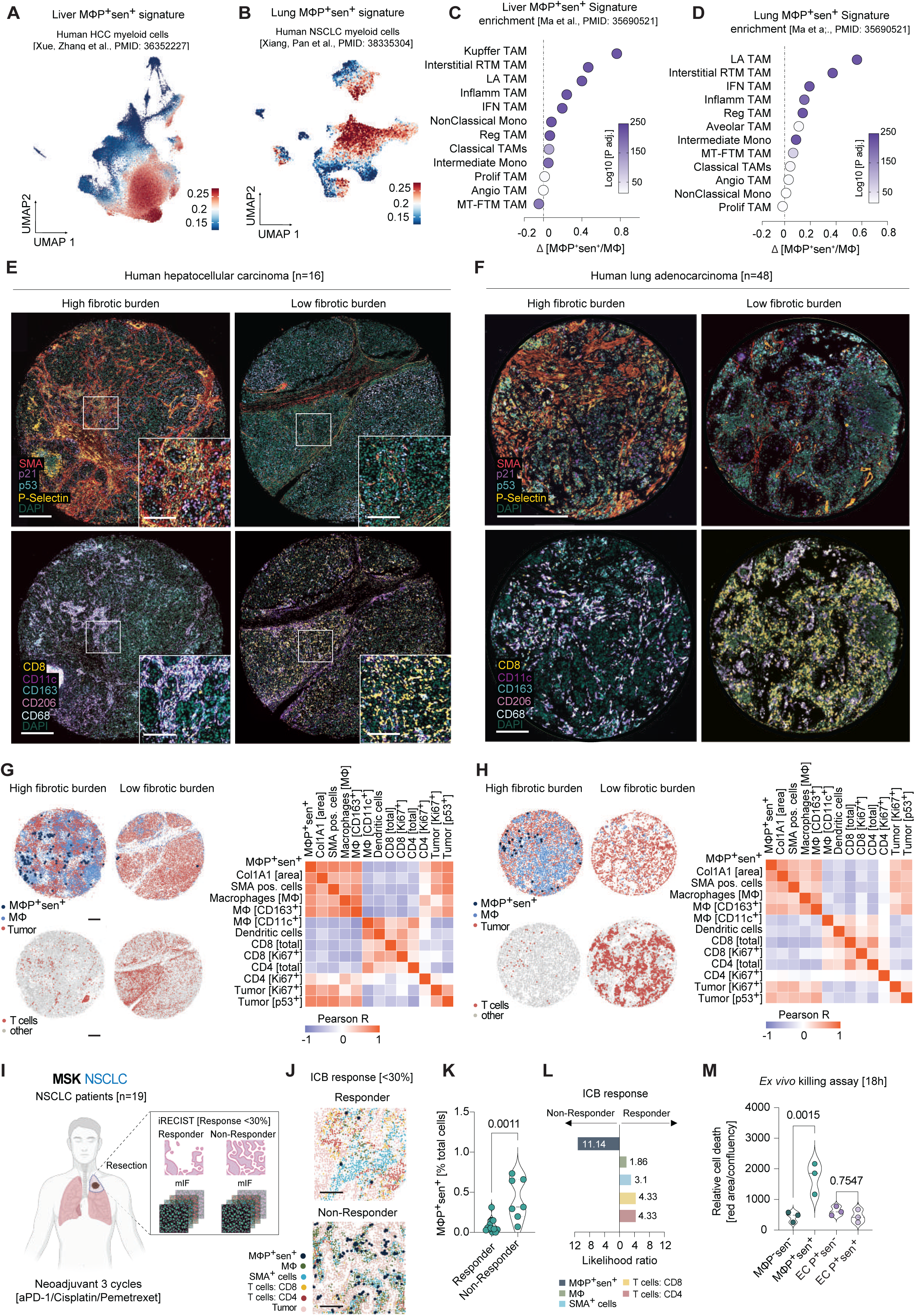
P-selectin⁺ senescent-like macrophages mark immunosuppressive, ICB-refractory human tumors and are vulnerable to SMNPs. (**A-D**) Enrichment of the P^+^sen^+^ macrophage (MΦP^+^sen^+^) gene signature in myeloid cells from human hepatocellular carcinomas (HCC) (A) or human non-small cell lung cancer (NSCLC) (B), and previously described tumor-associated macrophage (TAM) subsets from human HCC (C) or human NSCLC (D). (**E-F**) Representative mIF images of SMA, p21, p53, and P-selectin (top), and immune markers (bottom) on tissue microarrays (TMAs) of human HCC (E) and lung adenocarcinoma (LUAD) (F). Fibrotic burden was defined based on SMA expression. Scale bars, 500 μm. (**G-H**) (Left) Representative images from mIF analysis mapping MΦP^+^sen^+^, all macrophages (MΦ), and tumor cells comparing high and low fibrotic human HCC (G) and LUAD (H) TMAs. (Right) Cellular neighborhood analysis measuring the distance between the indicated immune and stromal populations in human HCC (16 TMAs) (G) and LUAD (48 TMAs) (H). (**I**) Schematic of human NSCLC biopsy. (**J**) Representative images from mIF analysis mapping MΦP^+^sen^+^, all macrophages, and tumor cells from NSCLC biopsies from responder and non-responder patients treated with immune checkpoint blockade (ICB). Scale bars, 250 μm. (**K**) Quantification of P-selectin^+^ senescent (P^+^sen^+^) macrophages in lung cancer of responder and non-responder patients to ICB (7-12 patients/group). (**L**) Likelihood ratio of presence of the indicated immune populations in lung cancers from ICB responder and non-responder patients. (**M**) Ex-vivo killing of sorted P^+^sen^+^ or P^+^sen^-^ MΦ and endothelial cells (EC) from human NSCLC biopsy treated with FiNav nanoparticles (3 human NSCLC patients). Each dot represents one patient. Data are presented as mean ± s.e.m. Statistical tests were Pearson correlation (G,H), Mann-Whitney test (K), likelihood ratio with cut-off defined using Youden’s J statistic based on ROC (L), and one-way ANOVA and Sidak’s multiple comparisons (M).

We extended our analyses to a broader tissue contact by performing mIF on 16 HCC and 48 NSCLC patients, where MΦP^+^sen^+^ cells comprised ∼2% and ∼0.4% of all cells, respectively (fig. S14E). Across both tumor types, higher MΦP^+^sen^+^ abundance was associated with highly proliferative, p53-mutated tumors (Ki67^+^ p53^+^ tumor cells) and fibrotic TMEs characterized by increased collagen deposition, αSMA^+^ fibroblasts, and macrophage infiltration (Fig. 6E-H). MΦP^+^sen^+^ abundance was positively correlated with CD163^+^ macrophages and inversely correlated with the presence of CD11c^+^ APC-like macrophages, dendritic cells, CD4^+^ cells, CD8^+^ cells, and proliferation (Ki67 low) (Fig. 6E-H). Spatial neighborhood analysis of highly fibrotic tumors mirrored murine findings, placing MΦP^+^sen^+^ macrophages within multicellular niches enriched for stromal activation and immune exclusion (fig. S14F). In a prospective cohort of 19 NSCLC patients receiving neoadjuvant anti–PD-1 plus carboplatin and pemetrexed (Fig. 6I), mIF revealed that MΦP⁺sen⁺ macrophages were significantly enriched in non-responding tumors (Fig. 6J, K), whereas intratumoral CD4⁺ and CD8⁺ T-cell abundance correlated with response, as expected (Fig. 6L).

To functionally probe SMNP activity in a human context, we obtained freshly resected NSCLC specimens from three patients, isolated macrophage and endothelial populations based on SPiDER activity, P-selectin expression, and lineage markers, and independently confirmed MΦP⁺sen⁺ macrophages by mIF (fig. S14G). Ex vivo treatment with FiNav, but not DexNav, selectively induced cell death in MΦP⁺sen⁺ macrophages with minimal effects on other macrophage subsets or endothelial cells (Fig. 6M).

To place these cells in a broader disease context, we applied the humanized MΦP⁺sen⁺ signature to a pan-tissue myeloid scRNA-seq atlas (*90*). The signature was highest in HES1⁺ macrophages (fig. S14H, I), an M2-like state enriched in osteoarthritis and rheumatoid arthritis (*90*). Furthermore, the humanized MΦP⁺sen⁺ signature enriched in breast, lung, liver and colon cancers (fig. S14J). Thus, MΦP⁺sen⁺ macrophages are conserved across human cancers and chronic inflammatory diseases, occupy pro-fibrotic, immune-excluded niches, predict immunotherapy resistance, and are selectively eliminated by SMNPs, thereby defining a tractable therapeutic axis linking senescent-like macrophage states to fibrosis-driven immune suppression.

## Discussion

Here we develop senescence-modulating nanoparticles (SMNPs) as a platform for selectively reprogramming fibrotic and immune-excluded tissue environments present in chronic inflammation and cancer. By coupling the high-affinity interaction between fucoidan and P-selectin to senolytic or senomorphic payloads, SMNPs concentrate senescence-modulating drugs within disease-relevant tissue niches. In murine models of liver and lung fibrosis, SMNPs reduced fibrotic burden, remodeled the tissue environment, and improved organ function. In tumor models embedded in fibrotic tissue, the same strategy remodeled stromal and myeloid barriers, restored immune infiltration, and reversed resistance to immune checkpoint-based therapies. This biological activity is coupled to a marked improvement in therapeutic index. SMNP delivery largely dissociated the senescence-modulatory activity of Navitoclax and BET inhibitors from the severe hematologic and systemic toxicities that constrained their further clinical development (*38, 39, 63–67*). This approach addresses two fundamental barriers to senotherapeutic translation: the inability of existing agents to discriminate pathogenic from reparative senescent-like populations and the systemic toxicity imposed by broad-acting agents.

The therapeutic selectivity of SMNPs reflects the convergence of three features. First, P-selectin provides a delivery handle, as fucoidan nanoparticles preferentially accumulate in P-selectin-positive cells and lose this advantage with dextran substitution or P-selectin blockade. Second, productive delivery is shaped by intracellular routing: P-selectin-positive endothelial cells engage fucoidan nanoparticles but route them through caveolin-associated transcytotic pathways (*46–48*), whereas P-selectin-positive senescent-like macrophages (MΦP⁺sen⁺) accumulate particles in endolysosomal compartments compatible with payload release. Third, the payload determines which P-selectin-positive macrophages are functionally affected. Fi-Navitoclax selectively eliminated the MΦP⁺sen⁺ population. However, Fi-Paclitaxel killed P-selectin-positive macrophages irrespective of senescence state but produced less antifibrotic benefit and greater toxicity. The comparable activity of dBET6-loaded SMNPs further indicates that suppressing senescence-associated secretory programs in specific pathogenic populations can be sufficient for therapeutic benefit. Thus, SMNP activity is not explained by P-selectin targeting alone, but by the intersection of targeted delivery, cell-type-specific nanoparticle processing, and payload activity matched to senescence-associated vulnerabilities.

Our study reinforces previous work indicating that the pathogenic effects of senescent cells on driving fibrosis is concentrated within discrete sub-populations rather than distributed uniformly across all cells bearing senescence markers (*16, 36*). Integrated single-cell, spatial, and functional analyses identify the MΦP⁺sen⁺ population as a key functional target of SMNPs in vivo. This population shares features with previously described pathogenic p16⁺ macrophages (*36*) and LAM states implicated in chronic inflammation, fibrosis, and immune suppression (*29, 36, 70, 75–77, 79*), but is not synonymous with all senescent-like macrophages or all LAMs. Rather, P-selectin marks a distinct senescent-like subset within the broader fibrosis-expanded LAM compartment, one in which senescence-associated programs, lipid handling, matrix remodeling, and immune suppression converge. This distinction is therapeutically important: SMNPs do not indiscriminately deplete all macrophages or even all LAMs but selectively engage in a disease-driving macrophage state conserved across murine models, human fibrotic tissues, and human tumors.

The cancer studies extend this concept from fibrosis resolution to immune restoration. In both liver and lung cancer models, the fibrotic microenvironment converted otherwise ICB-sensitive tumors into resistant tumors, directly modeling resistance patterns observed in patients with fibrotic tumor microenvironments (*10, 83*). SMNP treatment dismantled stromal and myeloid barriers, increased dendritic cell and T cell infiltration, and restored checkpoint blockade responsiveness to levels approaching those observed in non-fibrotic organs. The combination of FiNav and immune checkpoint blockade generated organized lymphoid aggregates, shifted dendritic cells toward mature and activated states, and required CD8^+^ and CD4^+^ T cells for therapeutic benefit. Human tumor analyses further supported clinical relevance of this axis. The MΦP⁺sen⁺ population were present in liver and lung cancers, enriched in fibrotic and immune-excluded niches, correlated with poor immunotherapy response, and are selectively vulnerable to FiNav ex vivo. These findings connect fibrosis-associated senescent-like macrophages to immune exclusion and identify their targeting to restore antitumor immunity. Moreover, the MΦP⁺sen⁺ transcriptional signature maps onto macrophage populations enriched across chronic inflammatory diseases including osteoarthritis and rheumatoid arthritis, suggesting that this pathogenic macrophage axis extends beyond cancer to a broader range of fibrosis-associated pathologies.

More broadly, these findings reinforce the view that senescence is not a monolithic cellular program but a heterogeneous spectrum of cell states with distinct, and sometimes opposing, roles in tissue repair, fibrosis, immune regulation, and cancer (*16, 18, 50, 91*). Although disease-driving macrophage states contribute to diverse pathologies (*29, 36, 70, 92–94*), the ability to selectively target these populations while sparing beneficial and reparative myeloid cells has remained a challenge. SMNPs address this gap by combining state-selective targeting with modular, niche-directed drug delivery, overcoming both biological and toxicity barriers that limit existing senotherapeutic approaches. More broadly, SMNPs establish a general principle for niche-directed therapy: therapeutic efficacy can be achieved by engaging small, pathogenic cell populations rather than broadly exposing entire tissues to drug. By combining interchangeable payloads matched to a conserved cellular target with clinically identifiable patient populations, SMNPs establish a translational framework for treating fibrotic disease and restoring immunotherapy responses in otherwise refractory cancers.

## Acknowledgments

We thank A. Gutierrez, E. Sisso and the MSKCC animal facility for technical support with animal colonies; the Molecular Cytology Core, the Center of Comparative Medicine & Pathology, the Genome Editing and Screening Core (RRID: SCR_027799), the Flow Cytometry Core (RRID: SCR_021105) and the Integrated Genomics Operation Core (RRID: SCR_027801) at MSKCC for assistance. We thank R. Xue and Z. Zhang for generating and sharing the scRNA-seq of liver cancer dataset via the National Genomics Data Center (GSA-Human). We thank A. Schietinger for advice and discussion. We thank Z. Liu and B. Zhou for sharing the p16^+^ macrophage signature from their liver fibrosis dataset, J. Novak for editing the manuscript; and all the members of the Lowe laboratory for advice and discussions. S.W.L. is an investigator in the HHMI and the Geoffrey Beene Chair for Cancer Biology. This article is subject to HHMI’s Open Access to Publications policy. HHMI laboratory heads have previously granted a nonexclusive CC BY 4.0 license to the public and a sublicensable license to HHMI in their research articles. Pursuant to those licenses, the author-accepted manuscript of this article can be made freely available under a CC BY 4.0 license immediately upon publication.

## Author contributions

C.H., V.J.A.B., and H.V.G. conceived the study, designed and performed experiments, analyzed data, and wrote the manuscript. K.C.V. and S.R. conceived the design of the fucoidan-dBET6 and fucoidan-navitoclax nanoparticles respectively. K.C.V., A.M.P., S.R., and L.R.H. produced nanoparticles. D.McH. and Y.-J.H. analyzed the RNA-seq data. K.C.V., A.M.P., S.R., L.R.H., A.C.-P., M.S., S.F., H.C.S., X.Z., G.D.B., X.L., J.T.W., W.L., J.S., T.T., R.G., and P.B.R. assisted with experiments, produced reagents, and/or discussed the data. N.R. and C.M.R. provided human lung tissue microarrays and discussed the data. M.J.B. provided human lung cancer samples and discussed the data. D.A.H. supervised the design and implementation of fucoidan nanoparticles. A.F. performed and supervised some of the experiments, analyzed data, and wrote the manuscript. S.W.L. and D.A.H. supervised experiments, provided funding, and wrote the manuscript. All authors read the manuscript.

## Data, code, and materials availability

The mouse scRNAseq datasets generated in this study have been deposited in the NCBI Gene Expression Omnibus (GEO) under accession number GSE319261. This study reanalyzes publicly available single-cell RNA-sequencing datasets: human liver and lung fibrosis (*79*), dataset available from the Broad Institute Single Cell Portal: https://singlecell.broadinstitute.org/single_cell/study/SCP2156/identification-of-a-broadly-fibrogenic-macrophage-subset-induced-by-type-3-inflammation-human-liver-lung-myeloid-fibrosis-scrnaseq-atlas; human liver cancer (*85*), GSA-Human Study HRA001748 (*36*), available via the National Genomics Data Center, human lung adenocarcinoma (*95*) available from the NCBI GEO (GSE253013), datasets on senescent tissue resident macrophages (*29, 70*) available from the NCBI GEO (GSE201247 and GSE203447). Due to their size, multiplex IF images cannot be easily deposited in a public repository and are available from the corresponding author (S.W.L) upon reasonable request.

## Material and Methods

### Patient samples

Human FFPE tissue microarrays (TMAs) were purchased from TissueArray (www.tissuearray.com). We used TMAs containing normal and cirrhotic human liver tissues (#LV805b), adjacent normal and fibrotic pulmonary tissue (#BC04118a), and human HCC (#LV487). Detailed clinicopathologic information for each case is available from the vendor. In addition, we used TMAs generated at Memorial Sloan Kettering Cancer Center (MSK): one comprising primary LUAD enriched in untreated patients (*96*), and the other comprising chemotherapy with or without immunotherapy treated LUAD patients. In addition, whole tissue sections and fresh human lung tumors specimens were obtained from MSK patients, all of whom consented to an institutional biospecimen banking protocol. All analyses were performed according to a biospecimen research protocol approved by the MSK institutional review board (IRB).

### Synthesis of Navitoclax, dBET6, and paclitaxel-encapsulated nanoparticles

Fucoidan nanoparticles were synthesized via methods adapted from (*97*). Nanoprecipitation was conducted by adding 450 µL of 10 mg/mL of fucoidan (Sigma-Aldrich, F8190) or dextran sulfate (Sigma-Aldrich, D8906), 100 µL of 0.01 mM sodium bicarbonate buffer (Fisher, S233; pH 8), and 100 µL of IR-125/indocyanine green (ICG; Thermo Scientific, 412541000, 2 mg/mL), mixed by vortexing. Fifty microliters of Navitoclax, dBET6, or paclitaxel dissolved in DMSO (20 mg/mL) was then added dropwise while vortexing. Nanoparticles were collected by centrifugation at 20,000g for 10 min for the dBET6 formulation and 30,000g for 10 min for Navitoclax and Paclitaxel formulations. Supernatants were removed and pellets were resuspended in 0.9% saline (Sigma-Aldrich, S8776) or water as appropriate. Drug loading was quantified using either the Alliance 2695e System and H-Class Ultra-Performace Liquid Chromatography (Waters) or the Agilent 1260 Infinity Bio-inert Quaternary LC (Agilent). Two-fold serial dilutions of drug were run on LC to generate standard curves, which were used to calculate the drug concentration. The nanoparticles were then diluted with 0.9% sodium chloride (Sigma, Cat #S8776) to a working concentration of 3.75 mg/mL.

### Microscopy for nanoparticle characterization

Atomic force microscopy (AFM) was used to visualize and characterize nanoparticle size and uniformity. Samples were deposited onto pretreated mica with 3-aminoproply-trietoxy silane (APTES) (Pelco Mica Disc, V1, Ted Pella) and left to air dry. AFM images were captured using the Nanowizard V (JPK-Bruker) in AC Mode Imaging at room temperature. AFM probe (FMV-A, Bruker) with nominal frequencies of approximately 75 kHz and nominal spring constant of 3N/m was used for imaging. Images were collected at a 1 Hz Line Rate with an image size of 1 x 1 μm to 3 x 3 μm at 512 x 512-pixel resolution. The images were processed with JPK Data Processing software.

### Bench characterization of Navitoclax, Paclitaxel and dBET6-encapsulated nanoparticles

To ensure batch consistency, nanoparticles were characterized for intensity-based hydrodynamic size, polydispersity index of the nanoparticles, and zeta potential in water using a Zetasizer Nano ZS (Malvern) and used only when they met quality control standards. Encapsulation efficiency was quantified by Ultra-Performance Liquid Chromatography (UPLC) using an Acquity C18 column (50 mm x 2.1 mm internal diameter; 1.7 μm; Waters Corporation) or by High-Performance Liquid Chromatography (HPLC) using C18 analytical columns (150 mm × 2.1 mm internal diameter, 3.5 μm; Agilent Technologies) and a mobile phase consisting of acetonitrile and deionized water, both containing 0.1% trifluoroacetic acid or formic acid. For UPLC, the program ran a gradient of 5% to 95% acetonitrile over 2 minutes, returning to 5% acetonitrile at 3.5 minutes with a flow rate of 0.5 mL/min. Retention times were 1.9 min for Navitoclax, 2.1 min for dBET6, and 2.2 min for Paclitaxel. For HPLC, the gradient elution program ran from 5% to 90% acetonitrile over 5 minutes, followed by 90% to 95% acetonitrile over 3 minutes, with a flow rate of 1 mL/min. Retention times were 5.3 minutes for Navitoclax and 5 minutes for dBET6, Paclitaxel was not analyzed in the Agilent HPLC system. Absorbance was monitored at 275 nm for Navitoclax and 256 nm for dBET6 for both UPLC and HPLC; Paclitaxel was monitored at 234 nm on UPLC. measured by dynamic light scattering (DLS) using the diluted samples.

### Nanoparticle Stability Assay

Navitoclax and dBET6 fucoidan nanoparticles were suspended in 100 µL of 0.9% saline and diluted 1:100 in water, Nanoparticle stability and size were measured by DLS from the diluted samples at different timepoints 0, 3, 6, 24, and 48 hours post nanoparticle formulation.

### Generation and authentication of *shSELP* cell lines

shRNAs targeting mouse *Selp* were generated as previously described. In brief, the SplashRNA algorithm (*98*) was used to generate 10 top-scoring shRNA, each of which was cloned into the miR-30 backbone (*99*). Renilla luciferase shRNA, previously described, was used as a control (*60, 100*). shRNA efficacy was tested by qPCR and western blot. The shRNA with the highest knock down efficacy (TGCTGTTGACAGTGAGCGCTCTCTGTAGTTTAAAACAAAATAGTGAAGCCACAGAT GTATTTTGTTTTAAACTACAGAGAATGCCTACTGCCTCGGA) was used in experiments. Expression of sh*Selp* was linked to a BFP reporter system, allowing for the sort and enrichment of sh*Selp*^+^ populations based on high BFP expression before experiments.

### Quantitative PCR with reverse transcription

Total RNA was isolated using the RNeasy Mini Kit (Qiagen) and cDNA was obtained using TaqMan reverse-transcription reagents (Applied Biosystems). Quantitative PCR (qPCR) was performed in triplicates using SYBR green PCR master mix (Applied Biosystems) on the ViiA 7 Real-Time PCR System (Invitrogen). *18S* (18S forward (5’-GTAACCCGTTGAACCCCATT-3’) and 18S reverse (5’-CCATCCAATCGGTAGTAGCG-3’) served as endogenous normalization controls for mouse samples.

### Animal Studies Housing conditions

All animal experiments were performed in accordance with protocols approved by the Memorial Sloan Kettering Cancer Center (MSKCC) Institutional Animal Care and Use Committee (protocol number 11-06-011, 12-04-006). MSKCC guidelines for the proper and humane use of animals in biomedical research were followed. Mice were housed in a temperature-controlled room (22 ± 1°C), with a 12-hr light/dark cycle and free access to food and water. All mouse experiments were performed on 8- to 14-week-old male mice of C57BL/6N background. Sample sizes were determined based on previous experiments and published studies to ensure adequate power to detect biologically relevant differences. Mice were randomly assigned to experimental groups. Investigators were blinded to group allocation during data collection and analysis whenever possible.

### In vivo induction of mouse liver and lung fibrosis models

To induce liver fibrosis, mice were treated twice a week with 12 consecutive intraperitoneal injections of 1 ml/kg carbon tetrachloride (CCl_4_) in peanut oil. Control mice received peanut oil (100 µL) alone on the same schedule. CCl_4_ treatment continued twice weekly upon nanoparticle administration. To induce lung fibrosis, bleomycin was administered intratracheally at 1.5 U/kg. with saline (50 µL) used as control. For nanoparticle treatment, mice received a weekly intravenous dose of 100 µL of Navitoclax-encapsulated (15 mg/kg), dBET6-encapsulated (25 mg/kg), or Paclitaxel-encapsulated (20 mg/kg) fucoidan nanoparticles, dextran control nanoparticles, or free fucoidan (1 mg/mL) for four weeks starting after liver fibrosis establishment (six weeks on CCl_4_) or once a week for three weeks starting after lung fibrosis establishment (7 days post-bleomycin administration). Samples were collected 4 days after the last injection.

### Fibrotic liver cancer model and treatment

Mice were first injected twice a week for 6 weeks with 1 ml/kg CCl_4_ to induce liver fibrosis (see above). Upon liver fibrosis establishment, 2.5 × 10^5^ murine liver cancer cells (*101*) (Myc-GFP; sg-Trp53; sh-Selp) were orthotopically transplanted into fibrotic livers. Upon tumor growth (2 weeks post-transplantation), mice were randomized into different groups based on tumor size, measured via ultrasound. Navitoclax-encapsulated (15 mg/kg) or dBET6-encapsulated (25 mg/kg) fucoidan (or dextran control) nanoparticles were injected intravenously at day 14, 17, 19 post transplantation then once a week until the end of the study. For first-line based ICB, anti-PD-L1 (BE0101, BioXCell, 5 mg/kg) and anti-VEGF-A (BE0399, BioXCell, 50 μg/mouse) depletion antibodies were administered intravenously at day 18 post transplantation twice or once a week, respectively. For immune cell depletion experiments, anti-CD4 (BE0003-1, BioXCell, 200 µg/mouse), anti-CD8 (BE0061, BioXCell, 200 µg/mouse), and anti-NK1.1 (BE0036, BioXCell, 250 µg/mouse) depletion antibodies were administered intravenously at day 18 post transplantation twice a week in combination with first-line ICB. As controls, rat IgG isotype (BE0090, BioXCell) depletion antibody was administered intravenously at day 18 post transplantation accordingly. Mice remained on CCl_4_ administration throughout the treatment period until collection and humane endpoint. Mice were monitored for liver tumor volume by ultrasound.

### Fibrotic lung cancer model and treatment

Mice were first injected once intratracheally with 1.5 U/kg bleomycin to induce lung fibrosis. Upon lung fibrosis establishment (day 7 post bleomycin injection), 1.5 × 10^5^ murine lung cancer cells (*102*) (KRas^G12D^; Trp53^KO^; sh-*Selp*) were injected intravenously. Upon tumor growth (1 week post transplantation), mice were randomized into different groups. Navitoclax-encapsulated (15 mg/kg) or dBET6-encapsulated (25 mg/kg) fucoidan (or dextran control) nanoparticles were injected intravenously at day 7, 10, 12 post cancer cell injection, then once a week for 3 weeks. For first-line based ICB, anti-PD-L1 depletion antibodies (BE0101, BioXCell, 5 mg/kg) and cisplatin (S1166, Selleck Chemicals, 3 mg/kg) were administered intravenously at day 18 post transplantation twice or once a week, respectively. For immune cell depletion experiments, anti-CD4 (BE0003-1, BioXCell, 200 µg/mouse), anti-CD8 (BE0061, BioXCell, 200 µg/mouse), and anti-NK1.1 (BE0036, BioXCell, 250 µg/mouse) depletion antibodies were administered intravenously at day 18 post transplantation twice a week in combination with first-line ICB. As controls, rat IgG isotype (BE0090, BioXCell) depletion antibody was administered intravenously at day 18 post transplantation accordingly. Mice were monitored for lung tumor volume by micro-computed tomography.

### Ultrasound and micro-computed tomography (µCT) imaging

For ultrasound, mice were first placed in a chamber for anaesthesia by isoflurane. Once mice were adequately anaesthetised, all abdominal hairs were removed using hair removal cream. The abdomen was cleaned off all excess cream with alcohol wipes to avoid skin irritation. Anaesthetised mice were then placed under constant nasal anaesthetic administration on a heated pad monitoring heart rate and breathing during the ultrasound scan. The shaved abdomen was covered in ultrasound gel before scanning to enhance the transducer detection. 2D scanning was then performed on the liver area to record the long (D) and short (d) tumor diameter using a Vevo F2 ultrasound (Fujifilm - VisualSonics). After scanning, mice were placed into an empty recovery cage and then transferred back to their original cage. Liver tumor volume was calculated using the following formula: tumor volume = (D x d^2^)/2. For µCT, scans were performed using a Mediso Nano SPECTCT system covering only the lung fields of each mouse as previously described (*103*).

### Bleeding/Coating time assessment

At the indicated time points and following the corresponding treatments, bleeding time was assessed by tail puncture. The duration from puncture to cessation of bleeding was recorded, with hemostasis confirmed by the absence of blood staining on a clean, dry gauze pad applied to the wound site.

### Animal tissue harvesting and blood/serum analysis

Mice were euthanized by carbon dioxide inhalation followed by cervical dislocation, and blood was collected by cardiac puncture. For lung collection, mice were systemically perfused with PBS to remove intravascular blood prior to harvest. Livers were excised and rinsed in PBS. The liver right lobe was divided into pieces for downstream processing as follows: 1. Formalin-fixed paraffin-embedded tissues (10% formalin (Epredia, 5705) for 24 hours then stored in 70% EtOH until paraffin embedding); 2. 4% paraformaldehyde (PFA)-fixed followed by cryopreservation. (4% PFA for 3 hours followed by fixation in 30% sucrose overnight before embedding in Tissue Plus O.C.T. Compounds (Fisher HealthCare, 4585) on dry ice); 3. flash freezing on dry ice for immediate use or storage at −80°C. Blood samples were processed to obtain EDTA-plasma and serum. Complete blood counts were performed using the Element HT5^TM^ hematology analyzer (Heska). Plasma and serum were isolated by centrifugation at 900g for 10 min at 4°C and stored at −80°C. Serum samples were submitted to the Laboratory of Comparative Pathology (Memorial Sloan Kettering Cancer Center [MSKCC]. For alanine aminotransferase (ALT) and aspartate aminotransferase (AST) enzymatic rate methods using automated analyzers were used.

### *In vivo* nanoparticle bioluminescence analysis

Upon tissue collection at indicated time point tissue was rinsed in PBS and kept on ice until imaging. Nanoparticle fluorescence in tissues was captured using iBright FL1000 (Invitrogen), 745em/765ex, and quantified using Fiji/ImageJ.

### Liver portal vein flow analysis

Mice were anesthetized and prepared for ultrasound as described before. The liver portal vein flow was recorded using the Pulse Wave Doppler function of the Vevo F2 ultrasound (Fujifilm - VisualSonics)

### Histological analysis: tissue preparation

Tissues for FFPE processing were fixed in 10% neutral buffered formalin (Epredia, 5705) for 24 h, transferred to 70% ethanol, and submitted to IDEXX or the Memorial Sloan Kettering Cancer Center (MSKCC) Molecular Cytology Core for paraffin embedding and sectioning. Hematoxylin and eosin (H&E) staining was performed using standard protocols. For immunohistochemistry (IHC) and immunofluorescence (IF) on FFPE tissues, blocks were sectioned at 5 μm, deparaffinized, and rehydrated through graded ethanol to water. Antigen retrieval was performed using a pressure cooker with either a dual–pH protocol (10 mM citrate buffer, pH 6.0 [Vector, H-3300], and EDTA–Tris buffer, pH 9.0 [Vector, H-3301]) or a single 10 mM citrate buffer, pH 6.0. For cryopreserved tissues, samples were fixed in 4% paraformaldehyde (PFA) in PBS (Thermo Fisher Scientific, J61899.AK) for 4–8 h, cryoprotected, and further fixed overnight in 30% sucrose in 4% PFA in PBS. Tissues were embedded in Tissue-Plus O.C.T. Compound (Fisher HealthCare, 4585) on dry ice and stored at −80°C. Cryosections (5–10 μm) were cut using a cryostat onto SuperFrost Plus microscope slides (Fisher Scientific, 12-550-15) and stored at −80°C until use. Prior to staining, slides were thawed at room temperature for ∼15 min or until condensation evaporated and rehydrated in PBS.

### Histological analysis: conventional immunohistochemistry and immunofluorescence

For formalin-fixed paraffin-embedded (FFPE) tissues, sections were deparaffinized, rehydrated, and heated to 95-97°C either in Antigen Unmasking Solution Citric Acid Based (pH 6.0) (Vector Laboratories, H-3300) or Tris-Based (pH 9.0) (Vector Laboratories, H-3301) for antigen retrieval. For immunohistochemistry assays, slides were washed with PBS and incubated with 3% hydrogen peroxide (Fisher scientific, H325-500) for endogenous peroxidase blocking. FFPE and cryosections tissue sections were outlined using an ImmEdge Hydrophobic Barrier PAP Pen (Vector Laboratories, H-4000) and blocked in PBS containing 3–5% BSA (GeminiBio, 700-100P) and 0.01–0.3% Tween-20 (PBST). Primary antibodies were diluted in blocking solution at the indicated concentrations and incubated on sections overnight at 4°C. Slides were washed in PBST and incubated with appropriate secondary antibodies diluted in blocking buffer for 1 h at room temperature or overnight at 4°C. Following washing, nuclei were counterstained with DAPI (1 μg/mL in PBS) for 5–10 min. Slides were then washed and coverslips mounted using ProLong™ Gold Antifade Mountant (Invitrogen, P36930). Slides were scanned using a Pannoramic Scanner (3DHistech) with a 20×/0.8 NA objective at the MSKCC Molecular Cytology Core, or images were acquired manually using a fluorescence microscope 20x,40x or 63x using an SP8 (Leica) or a Zeiss microscope (Imager.Z2, Zeiss camera (AxioCamMRc). For general quantification, 5 independent fields of views per mouse were used. Antibodies used are listed in Table S1.

### Sirius Red staining

Sirius red staining was performed using 5-um FFPE tissue section and the Sirius Red / Fast Green Collagen Staining Kit (Chondrex, 9046) following manufacturer’s instructions. Briefly, liver or lung tissue sections were baked, deparaffinized, and rehydrated through graded ethanol to water. Slides were stained with Dye solution for 30 minutes, washed and incubated with the dye extraction buffer. After rinsing in water, slides were dehydrated, cleared in xylene, and mounted with a resinous medium. Brightfield images were acquired under consistent illumination/white balance with Zeiss microscope (Imager.Z2, Hamamatsu camera (ORCA-R2)) at 20x magnification. Collagen area fraction (red) was quantified using ImageJ in 5 independent fields of views per mouse. Data were analyzed and visualized using GraphPad Prism.

### Detection of senescence-associated β-galactosidase activity

Chromogenic senescence-associated β-galactosidase activity (SAβ-gal) was performed on cryopreserved using X-gal (Invitrogen, 15520018) as previously reported (*104–106*). Briefly, tissues were washed twice with 1 mM MgCl2 PBS solution (pH 5.5 for mouse, pH 6.0 for human, adjusted using HCl). An X-gal working solution consisting of 1 mM MgCl2 PBS, 5 mM potassium ferricyanide, 5 mM potassium hexacyanoferrate (II), and X-gal (40X from 40 mg/mL stock diluted in N,N-dimethylformamide) was prepared, filtered (Steriflip Vacuum tube Top Filter Millipore, SE1M179M6) and applied to sections. Slides were incubated at 37°C in a humidified chamber for 2-6 hours. Upon signal detection, slides were washed in PBS 2X and coverslips were mounted using Fluoromount-G Mounting Medium (SouthernBiotech, 0100-01). Fluorescent detection of SA-βgal activity was carried out utilizing the ImaGene Red C_12_RG lacZ Gene Expression Kit (Invitrogen, I2906). Cryosections (5–10 μm) were incubated with 1% chloroquine at 37 °C for 30 min, washed twice with pre-cooled PBS, and incubated with C12RG (1:50 dilution) at 37 °C for 30 min to 2 h. Sections were rinsed immediately with PBS, where indicated, slides were subsequently processed for immunostaining following the protocol above. Fluorescence images were acquired as whole slide scans with a Pannoramic P250 Flash scanner (3DHistech, Hungary) using 20x/0.8NA objective lens by the Molecular Cytology Core. Fluorescent detection of SA-βgal activity following single-cell tissue dissociation as described below was performed using the SPiDER-βgal kit (Dojindo, SG04). Cells were resuspended in FACS buffer containing Baflomycin A1 1:1000 and incubatedfor 1 h at 37 °C followed by incubation with SPiDER-βgal substrate (1:1000) for 30 min at 37 °C. Cells were washed with FACS buffer and processed for flow cytometry and sorting as described below.

### Mutiplex IF imaging and analysis

Multiplex-IF experiments were performed using the cyclic IF CellDive platform (Leica) following the manufacturer’s protocol using either 5 µm FFPE or cryopreserved sections. Cryopreserved tissue sections were baked at 64C for 30 minutes and FFPE sections were prepared as above using a double pH6/pH9 antigen retrieval. Next all following steps were performed after securing the slides in the Leica Clickwell slide holder. Slides were washed with PBST and blocked with 3% BSA in PBST for 30 min to 1 h followed by counterstain DAPI for 5 to 10 min. After washes, slides were imaged unstained to acquire background autofluorescence. Samples were then sequentially stained with antibodies and imaged using DAPI, Cy3, Cy5, and FITC channels on the CellDive instrument with CellDive image acquisition and processing software. Each field of view (FOV) was imaged in each staining round, followed by autofluorescence removal, registration with baseline DAPI, and stitching. Unconjugated primary antibodies were used in the first round of staining, followed by secondary antibody staining. Antibodies used can be found in Table S1. After imaging, dye inactivation was performed using 0.1 M Na_2_CO_3_ 3% H2O2 solution for 15 minutes at RT before starting the next round of autofluorescence imaging and staining. All rounds of imaging and slide storage were done in a solution of PBS with 50% glycerol. Staining quality and fluorescence removal were verified after each round. The fully stitched images were imported into HALO image analysis software (Indica Labs) for analysis. Cell segmentation was performed using the “traditional” nuclear segmentation option, with analysis settings optimized for each staining category. Cell segmentation was performed using the “traditional” nuclear segmentation option, with analysis settings optimized for each staining category. Phenotypic classifications were used to define cell types (see Table S2 for details). When analyzing TMAs a limited number of cores or tissue areas were excluded from the analysis due to poor tissue quality, antibody aggregates, tissue folds or absence of tissue. Each patient’s data represents the average cell count across single or multiple replicate cores. After image processing, data were analyzed and visualized using GraphPad Prism. Spatial analyses were performed using the HALO image analysis software (Indica Labs). Distance between two cell populations was determined using the defined cell types (see above) using the nearest neighbor algorithm. To capture the spatial organization in situ where changes in both, cellular abundance and proximity might contribute to the biological phenotype, data presented here were not normalized to cell type abundance. Data were analyzed and visualized using GraphPad Prism.

### Cytokine profiling

After harvest, fresh liver or lung tissues were rinsed with PBS and placed in Tissue Lysis buffer (0.5% Igepal, 0.5% sodium deoxycholate, 0.1% sodium dodecyl sulfate, 50 mM Tris-HCl [pH 7.5], and 150 mM NaCl) with Protease Inhibitor Cocktail. Tissue was homogenized using mechanical homogenization and centrifuged at 1000g for 10 min at 4 °C. Supernatant was then quantified using BCA Protein kit following manufacturer instructions. The supernatant was used immediately or aliquoted and stored at −80 °C until use. Assays were using the Proteome Profiler Mouse XL Cytokine Array (R&D Systems, ARY022B) and the BioRad Imager (ChemiDoc MP Imaging System) with automated optimal exposure following the manufacturer’s protocols. To generate raw values, signal intensity was quantified using ImageJ. After image processing, data were analyzed and visualized using GraphPad Prism.

### Lung and liver dissociation

Single cell suspension of liver non-parenchymal cells was generated from normal or CCl_4-_induced fibrotic livers, 48 h after the final CCl_4_ or oil injection, using an in situ pronase-collagenase retrograde perfusion technique followed by Nycodenz density separation, as previously described (*107, 108*). Briefly, mice were anesthetized with isoflurane. The inferior vena cava (IVC) at the hepatic outflow was cannulated with a catheter IV (Terumo, SR-OX2225CA), the portal vein was cut to provide outflow, and the supra-diaphragmatic IVC was clamped to confine perfusion to the liver. Livers were perfused with pre-warmed PBS at 42 °C (5 ml/min), followed by pre-warmed pronase (0.4 mg/mL, Sigma, P5147) and finally by collagenase D (1.8 mg/mL, Roche, #11088874103) solutions. Perfused livers were excised and incubated ex vivo in pronase (0.5 mg/mL) and collagenase (0.4 mg/mL) solution for 30 min at 40 °C with gentle agitation, then mechanically dissociated and filtered through 70-μm strainers. Cell suspensions were twice washed in Gey’s balanced salt solution B, GBSS/B (Sigma-Aldrich, G9779) containing DNAase I (0.02 mg/mL, Roche, 10104159001). Cell suspensions were mixed 1:1 with 34% w/v Nycodenz to a final concentration of 17% w/v, overlaid with GBSS/B, and centrifuged at 1,380 × g for 17 min at 4°C with brake off; the interphase was collected. After washing, the final non-parenchymal cell pellets were resuspended in FACS buffer (PBS, 2 mM EDTA (Invitrogen 15575), 0.5% BSA (GeminiBio, 700-100P) for staining with SPIDER and fluorescent antibodies.

Dissociation of normal or fibrotic lungs (14 days after bleomycin or saline administration) was performed using a dispase-collagenase buffer that was used as previously described (*109*). Briefly, following systemic perfusion with PBS to clear lungs of blood, through the trachea lungs were inflated with 3 mL of lung digestion buffer (S-MEM (Thermo Fisher Scientific, 11380037) with 1.7 U ml−1 dispase (Corning, 354235), 500 U ml−1 collagenase IV (Thermo Fisher Scientific, 17104019) and 10 µg ml−1 DNase I (StemCell Technologies, 07900), finely minced and further dissociated in a 37°C oven with gentle agitation for 45 min. The dissociated cells were passed through a 100-µM filter and red blood cell lysis was performed using ACK lysis buffer (Thermo Fisher Scientific, A1049201), following the manufacturer’s protocol. Cells were washed with S-MEM containing 2% heat inactivated fetal bovine serum (HI-FBS; Hyclone, SH30910.03), filtered through a 40 µm strainer (Corning, 431750) and pelleted by centrifuging at 300g for 5 min at 4°C. Cell pellets were resuspended in FACS buffer for staining with SPIDER and fluorescent antibodies as described below.

Human lung tissue samples were obtained from 3 NSCLC patients undergoing pulmonary resections. Normal tissue was obtained outside of the tumor margin, determined by intraoperative pathology observation. Tissue samples were washed with pre-chilled PBS, minced, and centrifuged at 250g for 5 min at 4°C. The supernatant was drained and tissue samples were further minced and dissociated in 5 mL-10 mL of lung digestion buffer, described above. Samples were further dissociated in a 37°C oven with gentle agitation for 1-1.5hr. Following, dissociated tissue samples were processed as lung dissociation described above and resuspended in FACS buffer for downstream processing as described below.

### Flow cytometry

Following SPiDER staining when applicable, single-cell solutions of lung or liver cells were washed with pre-chilled PBS and stained with either LIVE/DEAD™ Fixable Near-IR Dead Cell Stain Kit (Liver digestions, Thermo Fisher Scientific, L34994) or LIVE/DEAD Fixable Aqua (Lung digestion, Thermo Fisher Scientific, L34957) per manufacturer’s instructions for 30 min on ice. Cells were then washed with PBS, then with FACS buffer and incubated blocked with anti-mouse Fc (BD Biosciences, 553142) in FACS buffer for 10 min on ice, followed by antibody panels for 30 min on ice. Stained cells were centrifuged at 250g for 5 min at 4°C, washed with FACS buffer and filtered into 5 mL polystyrene tubes with strainer Caps (Falcon, 352235). Single-cell suspension samples were acquired using a 5-laser Cytek Aurora spectral analyzer (RRID:SCR_019826) or sorted with a 5-laser BD FACSymphony S6 SE or BD FACSDiscover S8 (RRID:SCR_026674) cell sorters. 6-way sorting was performed using 4-way purity mode for the S6 and purity mode for the S8, with a 100 μm nozzle at 20 psi (137.9 kPa) and a drop drive frequency of 30 kHz. Spectral unmixing was done using vendor-specific acquisition software, with autofluorescence of non-senescent cells incorporated as an additional parameter. The resulting unmixed data were subsequently analyzed using FlowJo (RRID:SCR_008520).

For single cell RNA-sequencing sorts, cells were stained with SPiDER, BUV563-anti-CD45 (BD Biosciences, 612924, 1:200) and BB700-anti-CD62P (BD Biosciences, 742128, 1:200). Live/Dead^−^ singlets events were first split into CD45^−^ and CD45^+^ gates; within each gate, P-selectin^+^ (CD62P^+^) and P-selectin^−^ (CD62P^−^) as well as senescent (SPiDER^+^) and non-senescent cells (SPiDER^−^) subsets were defined and sorted simultaneously (see Fig. S4A, S5A for gating strategy). Because autofluorescence differs between CD45^−^ and CD45^+^ immune compartments, SPiDER negative thresholds were set separately in each CD45 gate, per the established method (*14*). For library preparation, CD45^+^ and CD45^−^ cells were pooled together into 4 main populations: SPiDER^−^/P-selectin^−^, SPiDER^+^/P-selectin^−^, SPiDER^−^/P-selectin^+^ and SPiDER^+^/P-selectin^+^ populations.

For sorts of lung and liver single cell suspensions used in *ex vivo* nanoparticle treatment experiments, panels included BUV563-anti-CD45 (BD Biosciences, 612924, 1:200), BB700-anti-CD62P (BD Biosciences, 742128, 1:200), APC-anti-CD31 (Biolegend, 102409, 1:200), and either PE-anti-CD64 (Biolegend, 139314, 1:200, for lung macrophages) or PE-anti-F4/80 (Invitrogen, 61-4801-82, 1:200, for liver macrophages). Endothelial cells were defined as Live/Dead^-^, CD45^-^, CD31^+^; macrophage as Live/Dead^−^, CD45^+^, CD64^+^ (lung) or Live/Dead^−^, CD45^+^, F4/80^+^ (liver) populations. Within each population SPiDER^−^/P-selectin^−^, SPiDER^+^/P-selectin^−^, SPiDER^−^/P-selectin^+^ and SPiDER^+^/P-selectin^+^ subsets were sorted and used for ex vivo nanoparticle treatment.

### Nanoparticle *Ex Vivo* Assays

Sorted cell populations, as described above, were centrifuged at 250g for 5 min at 4°C. Cell pellets were resuspended in DMEM (Gibco, 11995-065) supplemented with 10% FBS and 1% Penstrep (Gibco, 50-753-3040) containing either DexNav (1:200), FiNav (1:200), free Navitoclax (1:200), DexdBET6 (1:200), FidBET6 (1:200), DexPax (1:200), FiPax (1:200), or free Paclitaxel (1:200) and treated for 30 min. Cells were then centrifuged at 250g for 5 min at 4°C and washed 3 times with DMEM. Cells were resuspended in DMEM supplemented with 10% FBS and 1% PenStrep and seeded at 0.01×106 into a seeded in a 96-well plate (Cellvis, P96-1.5P), that was previously coated with collagen (30 min, 37°C, Advanced Biomatrix PureCol).

Following, to measure cell toxicity in samples treated with Nav or Pax loaded particles and their corresponding free drug, propidium Iodide (Thermo Fisher Scientific, P1304MP) was added to the milieu. After seeding the plate was loaded onto the IncuCyte S3 (Sartorius) and the Incucyte Software (v2020C) was used to quantify cell death and confluency over time.

In experiments using dBET6 particles, following seeding and treatment, the plate was placed in a humidified incubator at 37°C with 5% CO2 for 2hrs. After 2hrs, cells were fixed for 15 min on ice with 4% PFA in PBS. Subsequently, cells were blocked and permeabilized using 3 % BSA with 0.03 %Tween-20 for 30 min and then stained with anti-BRD4 (Abcam, ab128874, 1:200) in blocking buffer overnight at 4°C. After overnight incubation, cells were washed 3 times with blocking buffer and then incubated with Alexa Fluor 647-conjugated-anti-rabbit (Invitrogen, A32795, 1:1000) in blocking buffer for 1 h with gentle agitation. Cells were then washed with PBS and stained with DAPI (1 ug/mL) diluted in PBS for 5-10 minutes. After washes, cells were imaged using SP8 Leica microscope. Quantification of BRD4 signal was done using ImageJ software as described above.

### Single Cell RNA-sequencing library preparation

Cells were stained with Trypan blue and Countess II Automated Cell Counter (ThermoFisher) was used to assess both cell number and viability. Following QC, the single cell suspension was loaded onto Chromium GEM-X 3’ Chip (10x Genomics PN 1000690) and GEM generation, cDNA synthesis, cDNA amplification, and library preparation of 2-15,000 cells proceeded using the Chromium GEM-X Single Cell 3’ Kit v4 (10X Genomics PN 1000691) according to the manufacturer’s protocol. cDNA amplification included 11-12 cycles, and 88-327 ng of the material was used to prepare sequencing libraries with 10 cycles of PCR. Indexed libraries were pooled equimolar and sequenced on a NovaSeq 6000 (16588) or X (both projects) in a PE28/88 paired end run using the NovaSeq 6000 S4 or X 10B or 25B Reagent Kit (Illumina). An average of 26 thousand reads was generated per cell.

### Single Cell RNA-sequencing (scRNA-seq) data processing

#### Data preprocessing and quality control

All mouse scRNA-seq datasets were barcode-corrected, aligned and UMI-corrected using CellRanger (v8.0.1) using mouse genome mm10 and default parameters for 3’ v4 GEM-X scRNA-seq kit.

#### Empty droplet removal and ambient RNA subtraction

Empty droplets and ambient RNA subtraction were performed using the remove-background function of CellBender (v0.3.0) with *expected_cells* equal to estimated number of cells detected by CellRanger. *Total-droplets-included* was based on estimated UMI count plateau of ranked cell barcodes (Table S3), (*fpr* = 0.01 (default), *learning-rate* = 0.0001 (default) and *epochs* = 150 (default). Cellbender background-corrected count matrices were used for downstream analysis, and counts were summed for genes sharing the same gene symbol.

#### Low-quality cell removal, doublet detection & normalization

scRNA-seq data was processed using scanpy (v1.10.4) (*110*) and rapids_singlecell (0.10.10) (hereinafter rsc.). To remove low quality cells, quality control metrics for library size, percentage of mitochondrial (mt), ribosomal (ribo) and the number of genes with at least 1 count in a cell (cngeneson) were first calculated using (rsc.pp.calculate_qc_metrics). An additional ‘size_factor’ was calculated for downstream batch integration by dividing the total counts per cell (total_counts) by the average of the total counts for all cells in the dataset (hereinafter size_factor). Genes detected in fewer than 3 cells were filtered for downstream analysis. For quality control boundary criteria,loglibrary size (log1p_total_counts) < 7.5 and mitochondrial count percentage (pct_counts_mt) > 15 were used and cells falling below these criteria were excluded from downstream analysis. Doublets were removed using scrublet (rsc.pp.scrublet) with the following parameters (*log_transform*=True, *expected_doublet_rate*=0.1, *n_prin_comps*=30 (default), *threshold*=0.25, *sim_doublet_ratio*=2 (default)). Count data was normalized (rsc.pp.normalize_total – *target_sum* = 1×10^4^) and log transformed with pseudocount 1 (rsc.pp.log1p). This normalization was used for downstream processing with the exception of MAST differential expression analysis where *target_sum* = 1×10^6^.

#### Gene exclusion for post-cleaning preprocessing

The following classes of genes were excluded for downstream feature selection and embedding generation: (i) mitochondrial and (ii) ribosomal transcripts. In excluding these genes, we sought to minimize variation stemming from quality control metrics alone.

#### Embedding, batch integration and cell type annotation

To generate refined and batch corrected embeddings and batch corrected annotations across samples and conditions single-cell ANotation using Variational Inference (scANVI) was used (*111*). First a kNN graph was constructed on non-batch corrected count matrices. This was done by first selecting highly-variable genes (HVG) (rsc.pp.highly_variable_genes) with the following parameters: *flavor*=Seurat_v3, *n_top_genes*=3000, followed by dimensionality reduction using PCA (rsc.tl.pca) ((n_comp=500). A kNN graph was then generated using function rsc.pp.neighbors with *n_neighbors*=20 from the top 17 PCs as specified from the inflection point in the cumulative explained variance curve (explaining 58% of variance). To generate first approximate cell type annotations CellTypist (*112*) cell annotation (‘CellTypist_annot_voting’) was performed using celltypist.annotate function with *majority_voting* = True and using Immune_All_Highmodel. Immune_All_High model was converted from human to mouse using CellTypist’s convert function. Batch corrected latent space was calculated on CellBender corrected counts using only HVGs with the following model (scvi.model.SCVI.setup_anndata) parameters: *batch_key=’*SampleID’, *size_factor=’size_factor’, labels_key=* ‘CellTypist_annot_voting’ and (scvi.model.SCVI) with parameters: *n_layers*=2, *n_latent*=30. scVI latent space was then computed using scvi.model.train() function. Following, scANVI latent space was computed using scvi.model.SCANVI.from_scvi_model() and scvi.model.SCANVI.train(). kNN graph was then generated on scANVI latent space using rsc.pp.neighbors() with *n_neighbors* = *3*0. UMAP embeddings were generated with rsc.tl.umap() using default parameters. For downstream re-embedding of fibrotic liver/lung scRNA-seq datasets cell annotations were first re-generated utilizing batch corrected kNN graph and used to regenerate a scANVI latent space for liver and lung fibrotic samples with the same parameters used above. Final annotations for cell types were generated by first performing phenograph clustering using the phenograph package’s (v1.5.7) on scANVI latent space with function phenograph.cluster() with the following parameters (k=30, clustering_algo=’leiden’, resolution_parameter=0.8). Phenograph clusters were then annotated based on known markers for tissue cell type markers.

#### Differential expression

To perform single-cell differential expression 1×10^6^ normalized log-transformed gene counts was performed using MAST (v3.22) by first fitting hurdle model “∼1 + total_counts + SortID” with MAST zlm() function using parameters: *method*=’bayesglm’ and regularize=TRUE. Then likelihood ratio tests (LRT) were performed using MAST summary() function with *doLRT* corresponding to differential expression contrast of interest. FDR values from hurdle component < 0.05 and logFC > 0.5 or < −0.5 were used to determine differentially expressed genes. Differential expressed gene (DEG) analysis was used to generate gene signatures for MΦP^+^sen^+^ populations (compared to MΦP^-^sen^-^) in both the liver and lung.

### Analysis of single cell RNA sequencing data from fibrotic mouse models

We analyzed scRNA-seq data from mouse liver and lung fibrotic tissues to characterize the myeloid landscape during organ fibrosis. Preprocessed data (in .h5ad format) were integrated into Seurat for downstream analysis. Dimensionality reduction and visualization were performed using UMAP, with cell types annotated through multiple methods, including CellTypist and PhenoGraphclustering. Module scores for MΦP^+^sen^+^Liver and MΦP^+^sen^+^Lung signatures were calculated using the Seurat AddModuleScore function based on custom mouse-specific gene signatures. Comparative analysis of signature enrichment was conducted between specific sorted populations (e.g., *P^+^sen^+^* vs. *P*−*sen*−), with statistical significance assessed via Wilcoxon rank-sum tests and visualized through violin plots and feature plots to identify senescence-associated myeloid subsets across different fibrotic organs.

### Analysis of single cell RNA sequencing data from public datasets

#### scRNA-seq analysis of HCC patients

We analyzed scRNA-seq data of liver cancer patient samples from GSA-Human Study HRA001748 (National Genomics Data Center) (*85*). Specifically, we focused on the myeloid compartment of HCC patients, including macrophages, monocytes, and dendritic cells. To manage the computational scale of the dataset, count matrices were processed using BPCells. Data processing included standard Seurat normalization, cell cycle scoring, and regression of cell cycle effects and mitochondrial gene percentages. Integration across samples was performed using Harmony through Seurat’s IntegrateLayers function. Module scores were calculated for myeloid-specific signatures using the Seurat AddModuleScore function. Cells were classified into high/low groups based on the 75th percentile of MΦP^+^sen^+^Liver signature. We also used the study-defined TIMELASER annotations to contextualize our signature score across tumor immune microenvironment states and patient groups. Mean expression levels of senescence-related pathways within each clinical category. Statistical significance of signature score differences was assessed using Wilcoxon rank-sum tests, with results visualized through violin plots, box plots, and lollipop plots to display effect sizes and adjusted p-values.

#### scRNA-seq analysis in LUAD patients

We analyzed a previously published scRNA-seq dataset comprising 9 LUAD patient samples accessed through Gene Expression Omnibus (GEO; GSE253013), specifically focusing on tumor derived myeloid cell (including macrophages and monocytes) (*95*). To ensure computational efficiency for large-scale data, count matrices were managed using BPCells. Data processing included standard Seurat normalization, cell cycle scoring, and regression of cell cycle effects and mitochondrial gene percentages. Integration across samples was performed using Harmony through Seurat’s IntegrateLayersfunction. Module scores were calculated for senescence-associated and myeloid-specific signatures using the Seurat AddModuleScore function. Cells were categorized into high/low groups based on the 75th percentile of MΦP^+^sen^+^Lung gene signature. Statistical significance of signature score differences was assessed using Wilcoxon rank-sum tests, with results visualized through violin plots, box plots, and lollipop plots to illustrate median differences and adjusted p-values.

### scRNA-seq analysis in fibrotic and healthy tissues

We analyzed a previously published and pre-integrated scRNA-seq dataset of human liver and lung myeloid cells from healthy and fibrotic tissues (*79*). and available through the Broad Institute Single Cell Portal (https://singlecell.broadinstitute.org/single_cell/study/SCP2156/identification-of-a-broadly-fibrogenic-macrophage-subset-induced-by-type-3-inflammation-human-liver-lung-myeloid-fibrosis-scrnaseq-atlas). To facilitate the analysis of this large-scale dataset, count matrices were processed using BPCells for efficient storage and computation. Data processing included standard Seurat normalization, cell cycle scoring, and regression of cell cycle effects. Standardized embeddings from the original study were used for UMAP visualization. Module scores for senescence and progenitor signatures were calculated using Seurat’s AddModuleScore function with MΦP^+^sen^+^ combined signatures. This signature was generated by taking the overlap of genes with logFC of > 1 and FDR values of < 0.05 from the MΦP^+^sen^+^Liver and MΦP^+^sen^+^Lung signatures. To evaluate the enrichment of these signatures in disease states, cells were grouped based on health status (Healthy/Normal vs. Fibrosis), and statistical significance was determined using Wilcoxon rank-sum tests. Results were visualized through violin plots, feature plots, and average module score heatmaps to characterize the myeloid landscape across tissues. All signatures analyzed in this study are available in Table S4-6.

### Overlap Analysis of Differential Gene Signatures

Our mouse-derive gene signatures were compared against published senescent related signatures (*29, 36, 70*). (Overlapping genes were visualized as Venn diagrams by comparing upregulated or downregulated genes (logFC >1) in our mouse dataset (MΦP^+^sen^+^ vs MΦP^-^sen^-^) to the upregulated or downregulated genes in each publised dataset (MΦp16^+^ vs MΦp16^-^). Pathway enrichment analysis was performed on each gene partition using Enrichr (*72*) against Gene Ontology (GO) databases to identify functional annotations associated with specific transcriptional programs.

### Gene Signature Scoring in a Published Macrophage Atlas

To contextualize our mouse-derived gene signatures within a broader macrophage transcriptional landscape, gene signature scores were computed across the MoMac-VERSE atlas (*90*) using Seurat’s AddModuleScore function. Signature activity was visualized using FeaturePlot on the UMAP embedding and violin plots stratified by annotated cell type clusters. For each gene signature, additional horizontal violin plots were generated with tissues rank-ordered by median module score.

### Hazard ratio comparing High vs low gene signature scores

Overall survival (OS) info and TPM-normalized gene expression data were obtained from the UCSC Xena data portal. TPM values below −1 were excluded from downstream analysis. To evaluate the prognostic significance of our macrophage signatures, Singscore package (*113, 114*) was used to derive rank-based statistics. Kaplan-Meier survival curves were generated using the survminer R package, comparing survival outcomes between high and low expression groups stratified by median signature score levels. Statistical significance was assessed using the log-rank test, with p-values < 0.05 considered significant. The coxph function was used to fit the Cox proportional hazards model, with gene signature score as the primary variable and race and age included as covariates to account for potential confounders. Hazard ratios (HR) and 95% confidence intervals (CI) were calculated and visualized to determine the association between gene signature score and survival outcomes.

### Quantification and statistical analysis

For IHC/IF including multiplex IF studies, staining was analyzed using HALO (Indica labs) or Fiji softwares. For all *in vivo* studies, data are shown as means ± s.e.m.. Before performing any statistical tests, we tested normal distribution by Shapiro-Wilk test. Statistical significances were determined by two-tailed unpaired student’s or Welch’s t test for two-group comparison, Multiple group comparisons were performed using one-way ANOVA with Sidak’s multiple comparisons test. Overall survival (OS) was calculated using the Kaplan-Meier method and log-rank (Mantel-Cox) test was used for survival comparison. The predictive value of SMNP and MΦP⁺sen⁺ enrichment as a prognostic factor was evaluated by examining the area under the receiver-operator characteristic (ROC) curve using a confidence interval of 95%. Likelihood ratio was calculated using a cut-off defined via Youden’s J statistic based on Receiver operating characteristics (ROC) analysis. All statistical tests were considered statistically significant when P was less than 0.05. Statistical analysis was performed using GraphPad Prism (v10).

---

**Figure S1:**
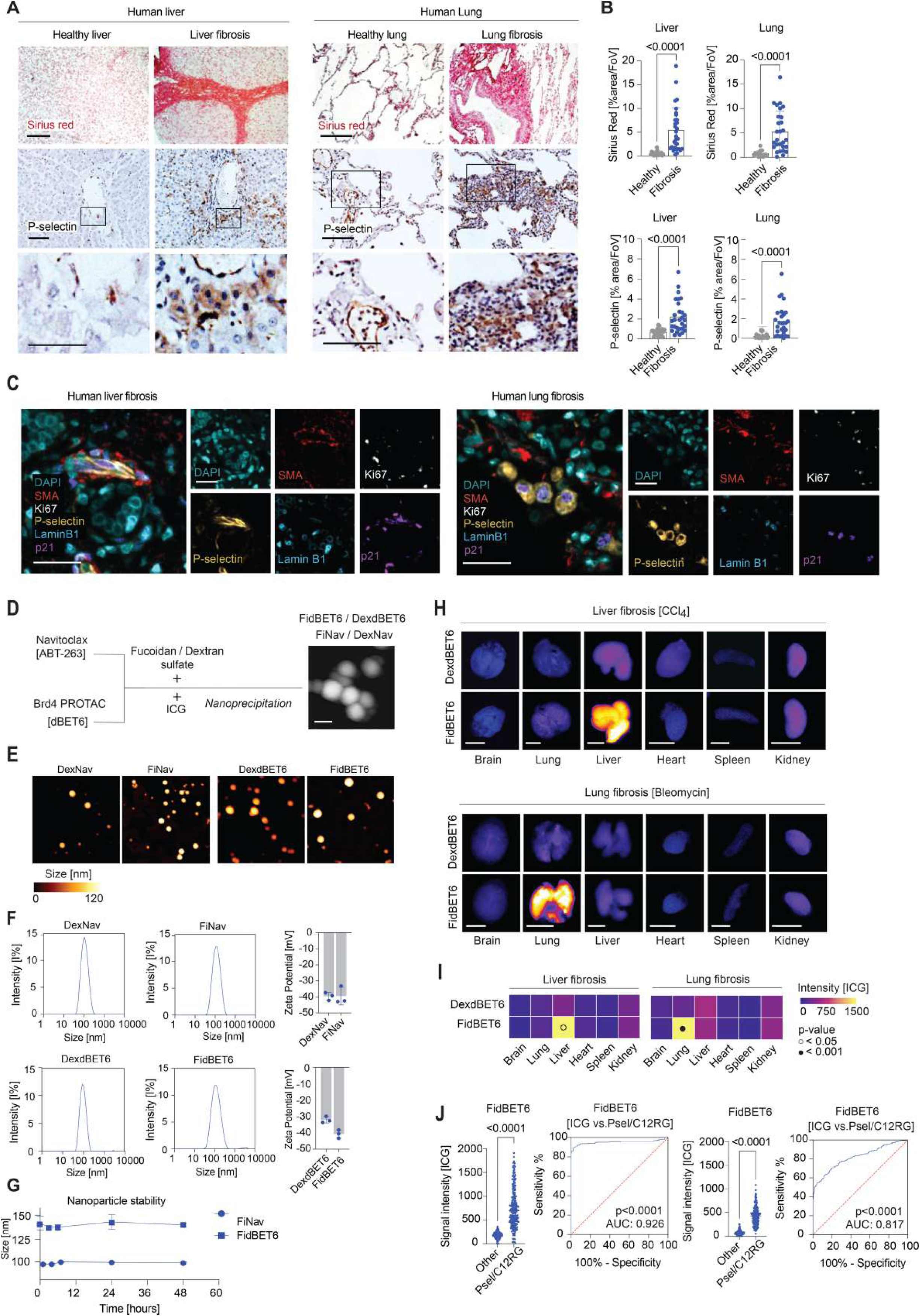
Expression and SMNP-mediated targeting of P-selectin-expressing cells in fibrotic tissues. (Related to Figure 1). (**A-B**) Representative images (A) and quantification (B) of collagen deposition (Sirius Red) and P-selectin expression (IHC) in human fibrotic and cirrhotic liver (healthy, n=21; fibrotic, n=30 cases) and lung (healthy, n=2; fibrotic, n=28 cases). Scale bars, 100 μm and 25 μm (magnification). (**C**) Representative images of P-selectin^+^ senescent cells, defined as p21^+^, Ki67^-^, LaminB1^low^ in human liver fibrosis (left) and lung fibrosis (right). Scale bars, 20 μm. (**D**) Schematic of drug (Navitoclax or dBET6) nanoprecipitation into sulfated fucoidan (Fi) or dextran (Dex) polysaccharides co-formulated with indocyanine green (ICG). (**E**) Representative atomic force microscopy images used to assess nanoparticle size (DexNav, FiNav, DexdBET6, FidBET6). (**F**) Dynamic light scattering (DLS) size distribution from representative individual nanoprecipitations of DexNav, FiNav, DexdBET6, and FidBET6 nanoparticles (left, middle) and surface zeta potential confirming dextran and fucoidan surface functionalization (right). (n=3 nanoprecipitation samples, 1 experiment). (**G**) FiNav and FidBET6 nanoparticle stability assay in saline. (**H-I**) Representative macroscopic images (H) and quantification (I) of ICG^+^ nanoparticle enrichment (FidBET6 and DexdBET6) across organs in murine models of liver and lung fibrosis at 12 h post nanoparticle injection; (3 mice/group). Scale bars, 500 μm. (**J**) ICG signal intensity in P-selectin^+^ senescent cells (defined as C12RG^+^; n=3 mice per group, left) and ROC curve of ICG specificity for P-selectin^+^ senescent cells (n=3 mice per group, right). Each dot represents one patient (B), one independent nanoparticle batch generated (F), or one cell (J). Data are presented as mean ± s.e.m. Mann-Whitney test (B,C,J), one-way ANOVA with Sidak’s multiple comparisons test (I), and receiver operating characteristic (ROC) analysis (J).

**Figure S2:**
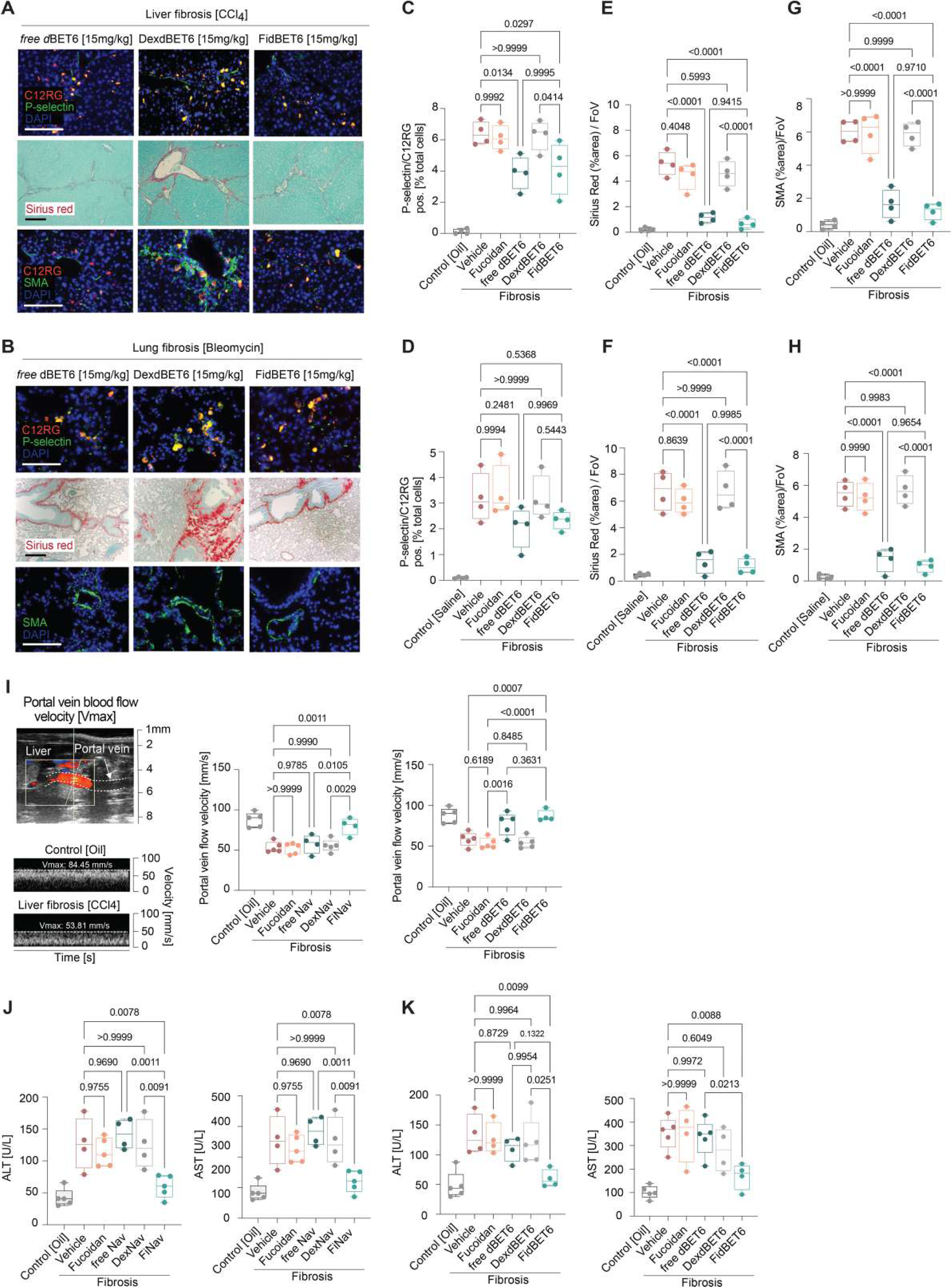
SMNPs improve fibrosis and organ function. (Related to Figure 2). (**A**-**H**) Analysis of collagen deposition (Sirius Red), fibroblast activation (SMA), and P-selectin^+^ senescent cells in CCl_4_-induced fibrotic livers (A) or bleomycin-induced fibrotic lungs (B) treated with free dBET6, Dex-dBET6 nanoparticles, or FidBET6 nanoparticles (A-B) and quantification (4 mice/group) including control groups (C**-**H). Scale bars, 150 μm. (**I**) Representative images (left) and quantification (right) of portal vein blood flow velocity in fibrotic livers treated as indicated (4-5 mice/group). (**J**-**K**) Analysis of alanine transaminase (ALT) (left) and aspartate transaminase (AST) (right) levels in CCl_4_-induced fibrotic livers treated with free fucoidan, (J) free navitoclax, DexNav, or FiNav; (K) free dBET6, DexdBET6, or FidBET6 (4-5 mice/group). Each dot represents one mouse. Data are presented as mean ± s.e.m. One-way ANOVA and Sidak’s multiple comparisons test used for all panels in this figure.

**Figure S3:**
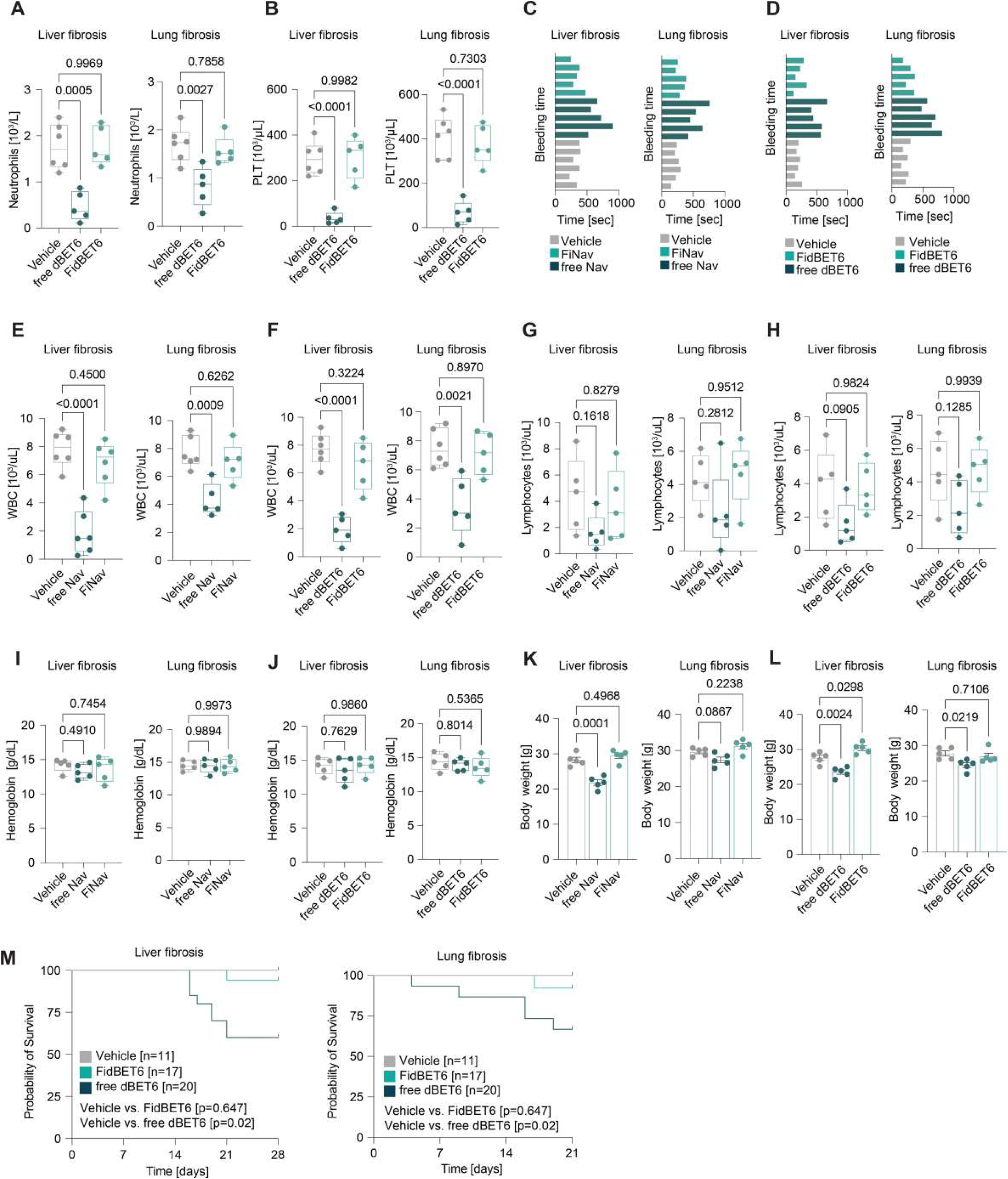
SMNPs eliminate drug-induced toxicities in mice with liver and lung fibrosis. (Related to Figure 2). (**A**) Neutrophil counts in the blood of mice with fibrotic livers (left) and lungs (right) treated with free dBET6 or FidBET6 nanoparticles. (**B**) Platelet counts in the blood of mice with fibrotic livers (left) and (lungs) treated with free dBET6 or FidBET6. (**C-D**) Bleeding time upon clotting after tail vein injection of (C) free navitoclax or FiNav nanoparticles; (D) free dBET6 or FidBET6 nanoparticles in fibrotic livers (left) and lungs (right). (**E-F**) White blood cell (WBC) counts in serum of mice with fibrotic livers (left) and (lungs) treated with (E) free navitoclax or FiNav; (F) free dBET6 or FidBET6. (**G-H**) Lymphocyte counts in serum of mice with fibrotic livers (left) and (lungs) treated with (G) free navitoclax or FiNav; (H) free dBET6 or FidBET6. (**I-J**) Hemoglobin counts in serum of mice with fibrotic livers (left) and (lungs) treated with (I) free navitoclax or FiNav; (J) free dBET6 or FidBET6. (**K-L**) Body weight of mice with fibrotic livers (left) and lungs (right) treated with (K) free navitoclax or FiNav; (L) free dBET6 or FidBET6. (**M**) Kaplan-Meier analysis comparing overall survival of mice with fibrotic livers (left) or fibrotic lungs (right) treated with free dBET6 or FidBET6. All data were collected after 4 weeks of treatment in fibrotic livers and 3 weeks of treatment in fibrotic lungs (5-6 mice/group). Each dot or bar represents one mouse. Data are presented as mean ± s.e.m. One-way ANOVA with Sidak’s multiple comparisons test used for (A-B, E-L), and log-rank Mantel-Cox test (M).

**Figure S4:**
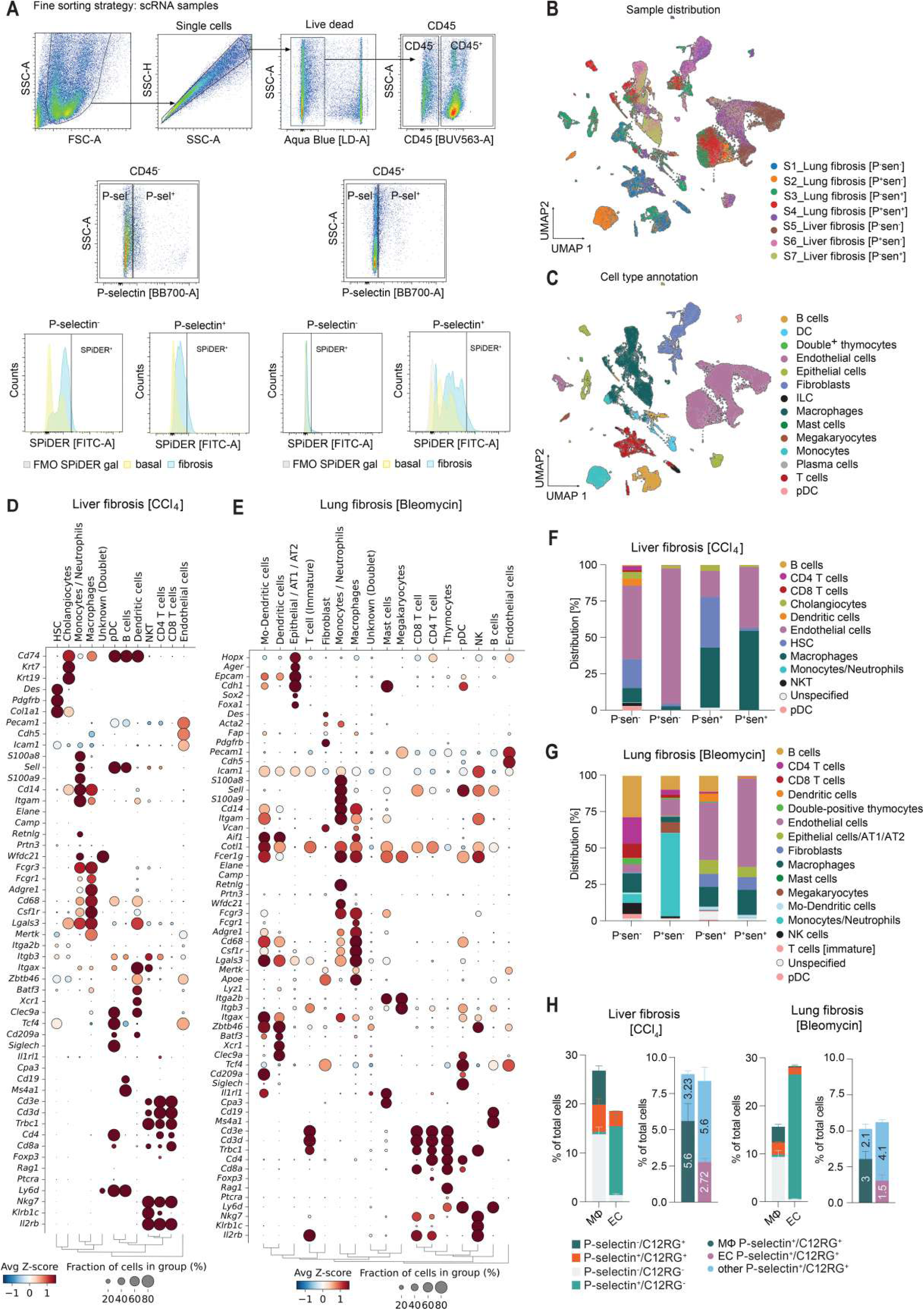
Characterization of P-selectin positive states in fibrotic liver and lungs. (Related to Figure 3). (**A**) Gating strategy for the isolation of live CD45^+^ and CD45^-^ populations based on P-selectin surface expression and b-galactosidase activity (SPiDER) in healthy and fibrotic tissues. (**B**) Visualization of unsupervised clustering in a UMAP of cells isolated from murine fibrotic liver and lung, sorted as in (A), then subjected to scRNA-seq. (**C**) Cell type annotations of the UMAP in (B). (**D-E**) Dot plots showing expression levels of representative marker genes within each cell population from fibrotic liver (D) and lung (E). (**F-G**) Proportions of the indicated cell types across sorted samples in fibrotic livers (F) and lungs (G). (**H**) (Left) Proportion of P-selectin^-^/C12RG^+^, P-selectin^+^/C12RG^+^, P-selectin^+^/C12RG^-^, and P-selectin^-^/C12RG^-^ macrophages (MΦ) and endothelial cells (EC) from sorted samples from fibrotic livers and lungs. (Right) Proportion of P-selectin^+^/C12RG^+^ MΦ, P-selectin^+^/C12RG^+^ EC, and other P-selectin^+^/C12RG^+^ cells in fibrotic livers and lungs (related to Fig. 3E-F).

**Figure S5:**
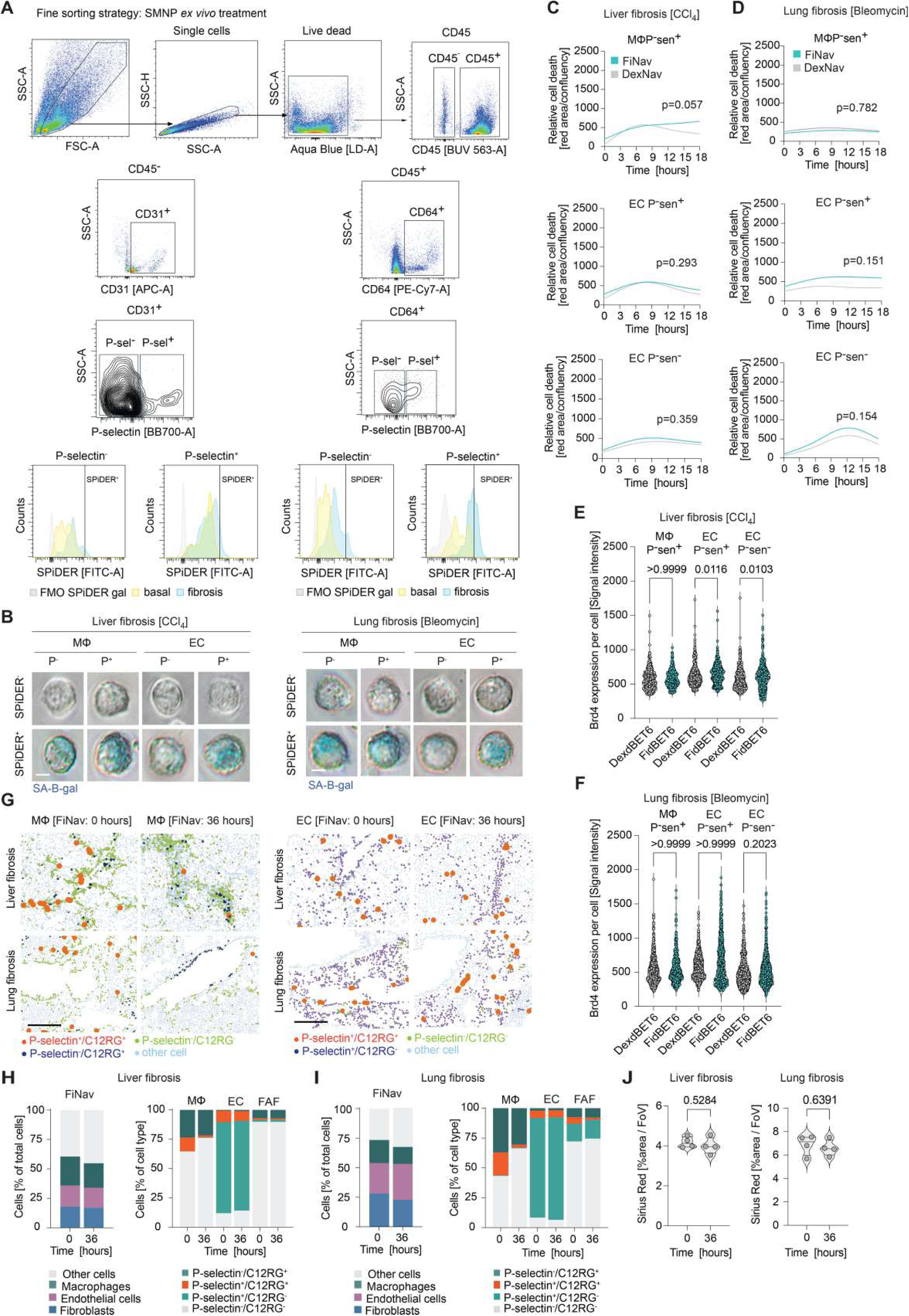
Functional assessment of SMNP target populations in fibrotic liver and lungs. (Related to Figure 3). (**A**) Gating strategy for the isolation of CD31^+^ endothelial cells (EC) (left) and CD64^+^ macrophages (MΦ) (right) based on their P-selectin expression and SA-b-galactosidase activity (SPiDER) in healthy and fibrotic murine lungs. (**B**) Representative images of senescence-associated β-galactosidase activity in the different sorted population of EC and MΦ from (A). Scale bars, 5 μm. (**C-D**) Ex-vivo killing assays of sorted P-selectin^-^ senescent^+^ (P-sen^+^) MΦ, P^-^ sen^+^ EC, or P-selectin^-^ senescent^-^ (P^-^sen^-^) EC from murine fibrotic livers (C) and lungs (D) treated with DexNav or FiNav nanoparticles. Representative data from three independent experiments. (**E-F**) Quantification of Brd4 protein after DexdBET6 or FidBET6 treatment of P^-^sen^+^MΦ, P^-^sen^+^ EC, and P^-^sen^-^ ECs from murine fibrotic livers (E) and lungs (F). Data are from four independent biological replicates and presented as mean ± s.e.m. (**G**) Representative images of mIF analysis of P-selectin^+^/C12RG^+^, P-selectin^-^/C12RG^+^, and P-selectin^-^/C12RG^-^ MΦ (left) and ECs (right) from fibrotic livers (E) and lungs (F) before (0 hour) and after (36 hours) FiNav injection. Scale bars, 250 μm. (**H-I**) Proportions of various cell types from fibrotic livers (H) and lungs (I) before (0 hour) and after (36 hours) FiNav injection. (Left) Macrophages, endothelial cells, and fibroblasts; (right) P-selectin^+^/C12RG^+^, P-selectin^-^/C12RG^+^, P-selectin^+^/C12RG^-^, and P-selectin^-^/C12RG^-^ MΦ, ECs, and fibroblasts (FAF) (4 mice/group). (**J**) Quantification of collagen deposition (Sirius Red) in fibrotic livers (left) and lungs (right) before (0 hour) and after (36 hours) FiNav injection (4 mice/group). Each dot represents one cell. Statistical tests were Mann-Whitney test or Welch’s t-test at endpoint (18h) for (C,D), Mann-Whitney test for (E,F), and unpaired t-test were used for (J).

**Figure S6:**
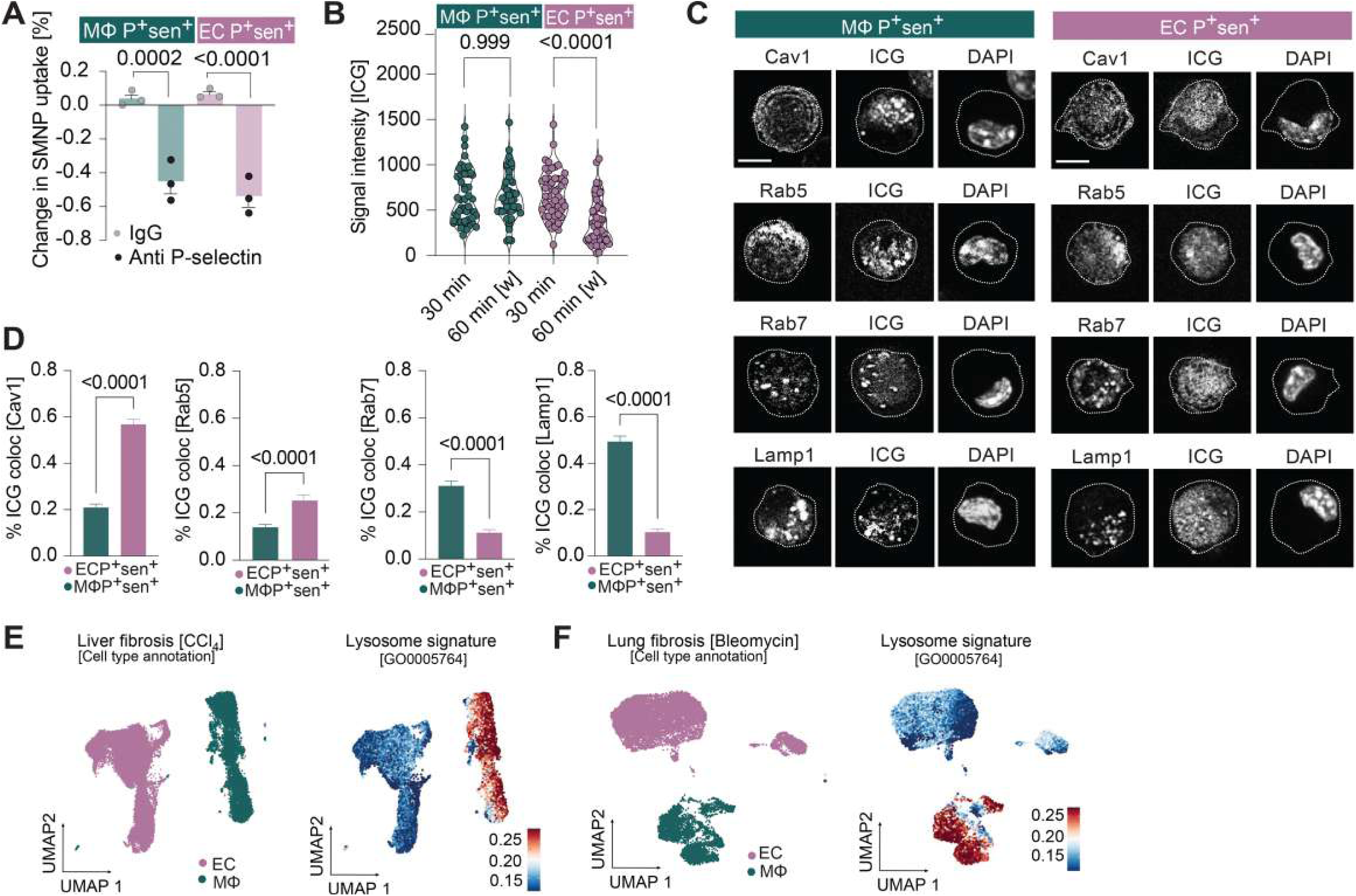
Characterization of uptake and intracellular trafficking of SMNPs. (Related to Figure 3). (**A**) SMNP (FidBET6) uptake in sorted P^+^sen^+^ macrophages (MΦ) and endothelial cells (EC) from murine fibrotic lungs following 2-hour treatment with FidBET6 nanoparticles in combination with P-selectin blocking antibody or IgG control. Change in NP uptake is relative to IgG controls. (**B**) Quantification of nanoparticle signal (ICG) in sorted P^+^sen^+^ EC and MΦ from murine fibrotic lungs after either a 30-minute exposure to FidBET6 nanoparticles or 30 minutes of exposure followed by a 30-minute washout (60 min [w]). (**C**) Representative IF images of nanoparticle signal (ICG) with vesicular compartment markers: Cav1 (transcytosis-associated marker), Rab5 (early endosomes), Rab7 (late endosomes), and Lamp1 (lysosomes) in sorted P^+^sen^+^ EC and MΦ from murine fibrotic lungs following treatment with FidBET6 nanoparticles. (**D**) Quantification of ICG^+^ SMNP colocalization with Cav1 (far left), Rab5 (left), Rab7 (right) or Lamp1 (far right) from images in C. (**E-F**) Lysosome signature in ECs and MΦ sorted from murine fibrotic liver (E) and lung (F). Left, cell type annotations in a UMAP of ECs and MΦ; right, enrichment of signatures for lysosomes. Each dot represents the mean signal across replicates (A), or one cell (B). (A and D) Data are presented as mean ± s.e.m. Mann Whitney test was used for (A-B, D).

**Figure S7:**
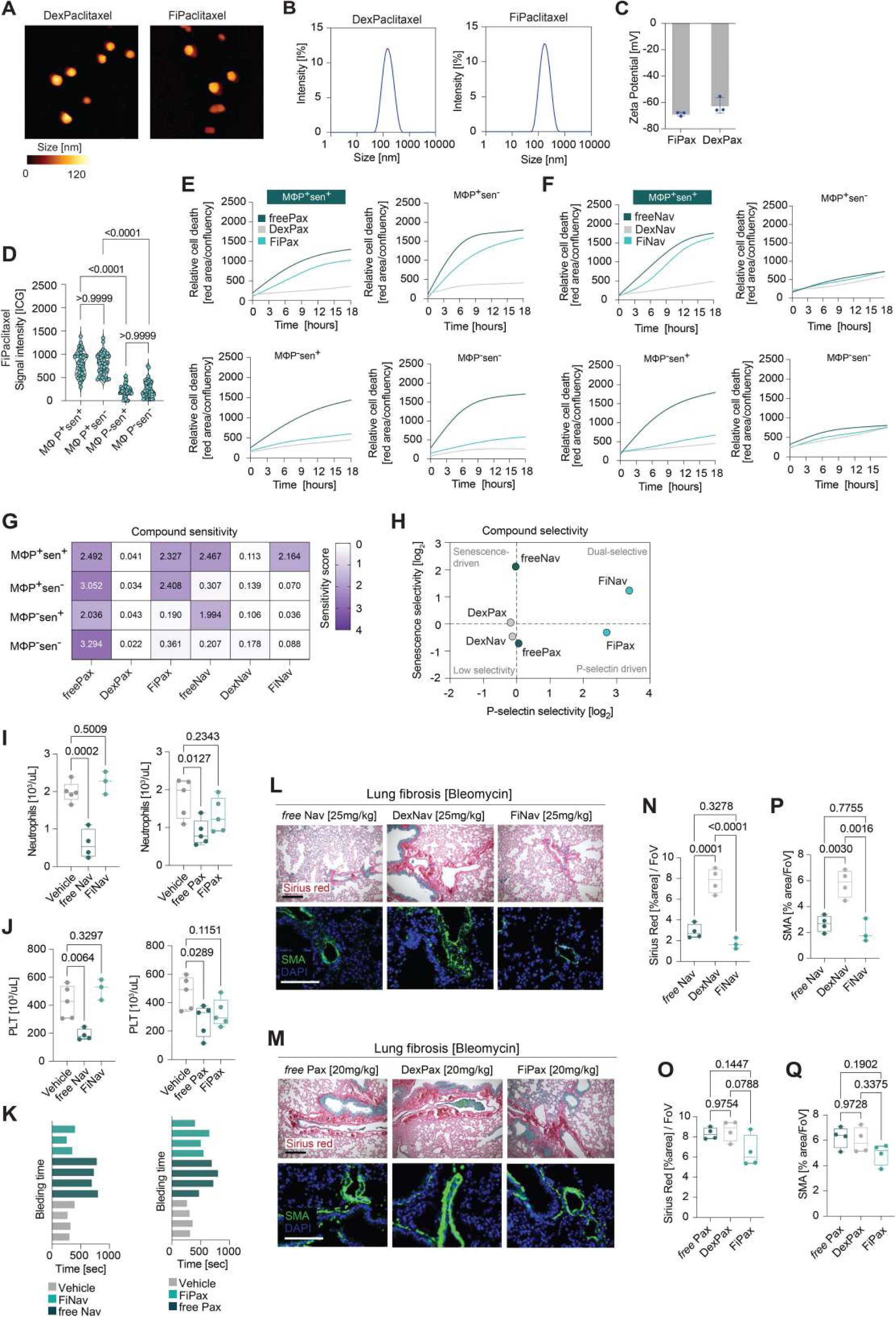
Characterization and activity of non-senotherapeutic nanoparticles. (Related to Figure 3). (**A**) Representative atomic force microscopy images used to assess the size of paclitaxel-encapsulating dextran nanoparticles (DexPax) and fucoidan nanoparticles (FiPax). (**B**) Dynamic light scattering size distribution from representative individual nanoprecipitations of DexPax (left) and FiPax (right). (**C**) Surface zeta potential confirming dextran and fucoidan surface functionalization (n=3 nanoprecipitation samples, 1 experiment). (**D**) Quantification of nanoparticles (ICG) in sorted P^+^sen^+^, P^+^sen^-^, P^-^sen^+^, and P^-^sen^-^ MΦ from murine fibrotic lungs following treatment with FiPax. (**E-F**) Ex-vivo killing of sorted P^+^sen^+^, P^+^sen^-^, P^-^sen^+^, and P^-^sen^-^ MΦs from murine fibrotic lungs treated with (E) free Paclitaxel (Pax), DexPax, or FiPax and (F) free Nav, DexNav, or FiNav. (**G**) Sensitivity table of freePax, DexPax, FiPax, freeNav, DexNav, and FiNav drugs for the indicated macrophage populations in fibrotic lungs. (**H**) Selectivity of free Pax, DexPax, FiPax, free Nav, DexNav, and FiNav for senescence, P-selectin, or both in fibrotic lungs. (**I-J**) Neutrophil (I) and platelet (J) counts from blood of mice with fibrotic lungs treated with (left) free navitoclax or FiNav and (right) free Paclitaxel or FiPax (3-5 mice/group). (**K**) Bleeding time upon clotting after tail vein injection of (left) free navitoclax or FiNav, and (right) free Paclitaxel or FiPax in fibrotic lungs. (**L-M**) Representative images of collagen deposition (Sirius Red), and fibroblast activation (SMA), in fibrotic lungs treated with (L) free Nav, DexNav, or FiNav or (M) free Pax, DexPax, or FiPax. Scale bars, 150 μm. (**N-O**) Quantification of collagen deposition (Sirius Red) in fibrotic lungs treated with (N) free Nav, DexNav, or FiNav or (O) free Pax, DexPax, or FiPax (3-4 mice/group). (**P-Q**) Quantification of myofibroblast activation in fibrotic lungs treated with (P) free Nav, DexNav, or FiNav or (Q) free Pax, DexPax, or FiPax (3-4 mice/group). Each dot represents one independent nanoparticle batch generated (C), one cell (D), or one mouse (I-J,N-Q). Each bar represents one mouse. (C-D, I-J, N-Q) Data are presented as mean ± s.e.m. One-way ANOVA with Sidak’s multiple comparisons test was used for (D, I-J, N-Q).

**Figure S8:**
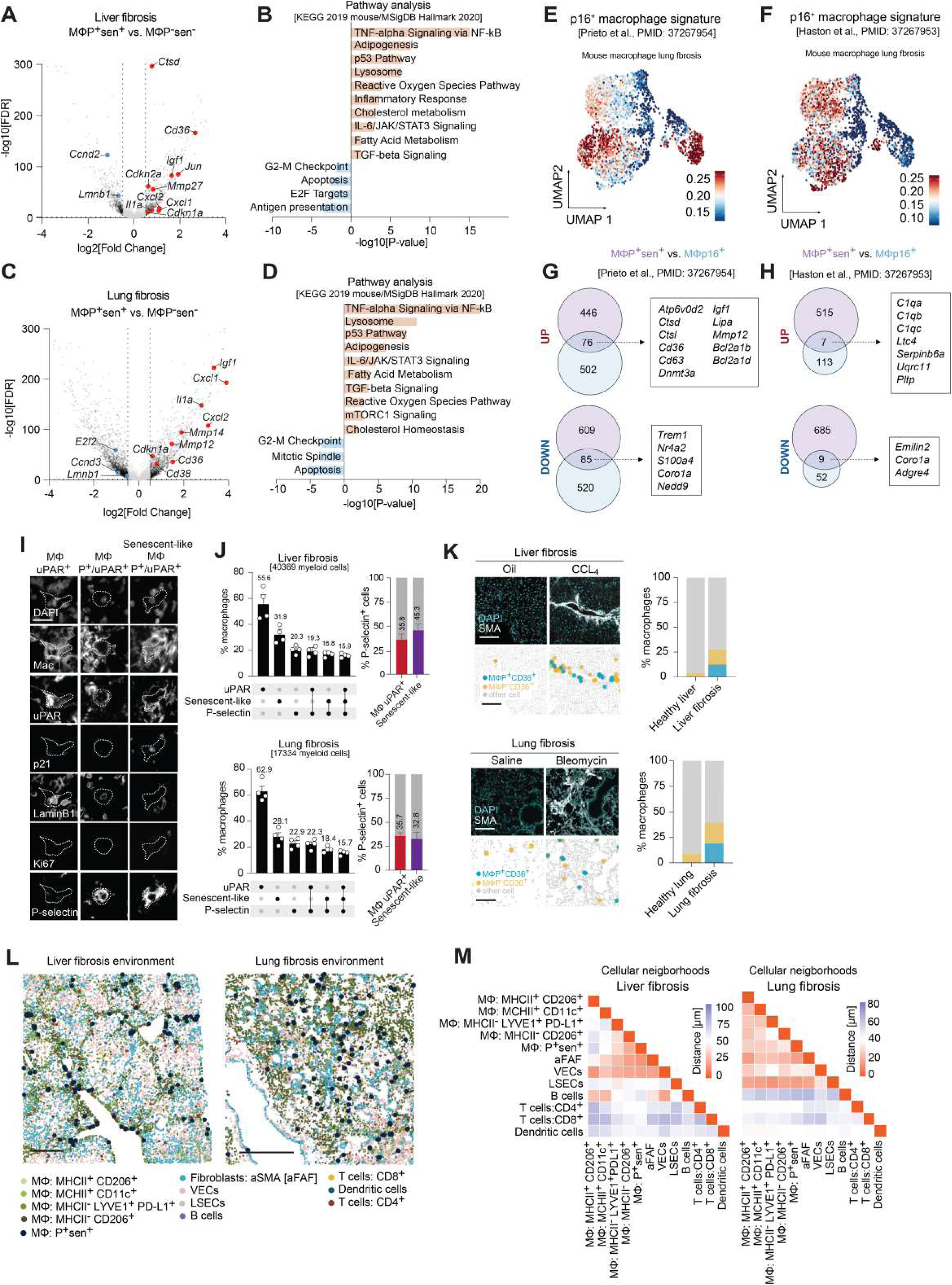
Transcriptional and spatial characterization of P+sen+ macrophages. (Related to Figure 4). (**A**) Volcano plot showing genes differentially expressed in P^+^sen^+^ macrophages compared to P^-^ sen^-^ macrophages in liver fibrosis. Representative senescence-associated genes are highlighted. (**B**) Upregulated and downregulated pathways of P^+^sen^+^ macrophages vs P^-^sen^-^ macrophages in murine fibrotic livers. (**C**) Volcano plot showing genes differentially expressed in P^+^sen^+^ macrophages compared to P^-^sen^-^ macrophages in lung fibrosis. Representative senescence-associated genes are highlighted. (**D**) Upregulated and downregulated pathways of P^+^sen^+^ macrophages vs P^-^sen^-^ macrophages in murine fibrotic lungs. (**E-F**) Gene signature analysis of P^+^sen^+^ macrophages compared to P^-^sen^-^ macrophages. The UMAP plots show enrichment of signatures for p16^+^ macrophages in murine lung fibrosis. (**G-H**) Venn diagrams representing the common upregulated (UP) or downregulated (DOWN) genes (derived from scRNA data) between P^+^sen^+^ macrophages compared to p16^+^ macrophages in murine lung fibrosis. (**I**) mIF analysis of uPAR, p21, LaminB1, Ki67, and P-selectin expression in macrophages from fibrotic murine livers. Senescent-like phenotype is defined by p21-positivity and absence of Ki67 and LaminB1 expression. (**J**) (Left) Quantification of uPAR, senescent-like, and/or P-selectin expression in macrophages and (right) quantification of uPAR and senescent-like expression in P-selectin^+^ myeloid cells of murine liver (top) and lung (bottom) fibrosis. (**K**) (Left) Representative images of mIF analysis mapping CD36^+^ and P-selectin^+^/CD36^+^ macrophages and (right) quantification of P^+^CD36^+^ and P^-^CD36^+^ macrophages among total macrophages of healthy and fibrotic murine liver (top) and lung (bottom). Scale bars, 250 μm. (**L**) Representative images of mIF analysis mapping different immune and stromal populations in murine fibrotic livers (left) and lungs (right). Scale bars, 250 μm. VEC: vascular endothelial cells; LSEC: Liver sinusoidal endothelial cells. (**M**) Cellular neighborhood analysis measuring the distance between the indicated immune and stromal populations in murine fibrotic livers (left) and lungs (right). Each dot represents one mouse (J).

**Figure S9:**
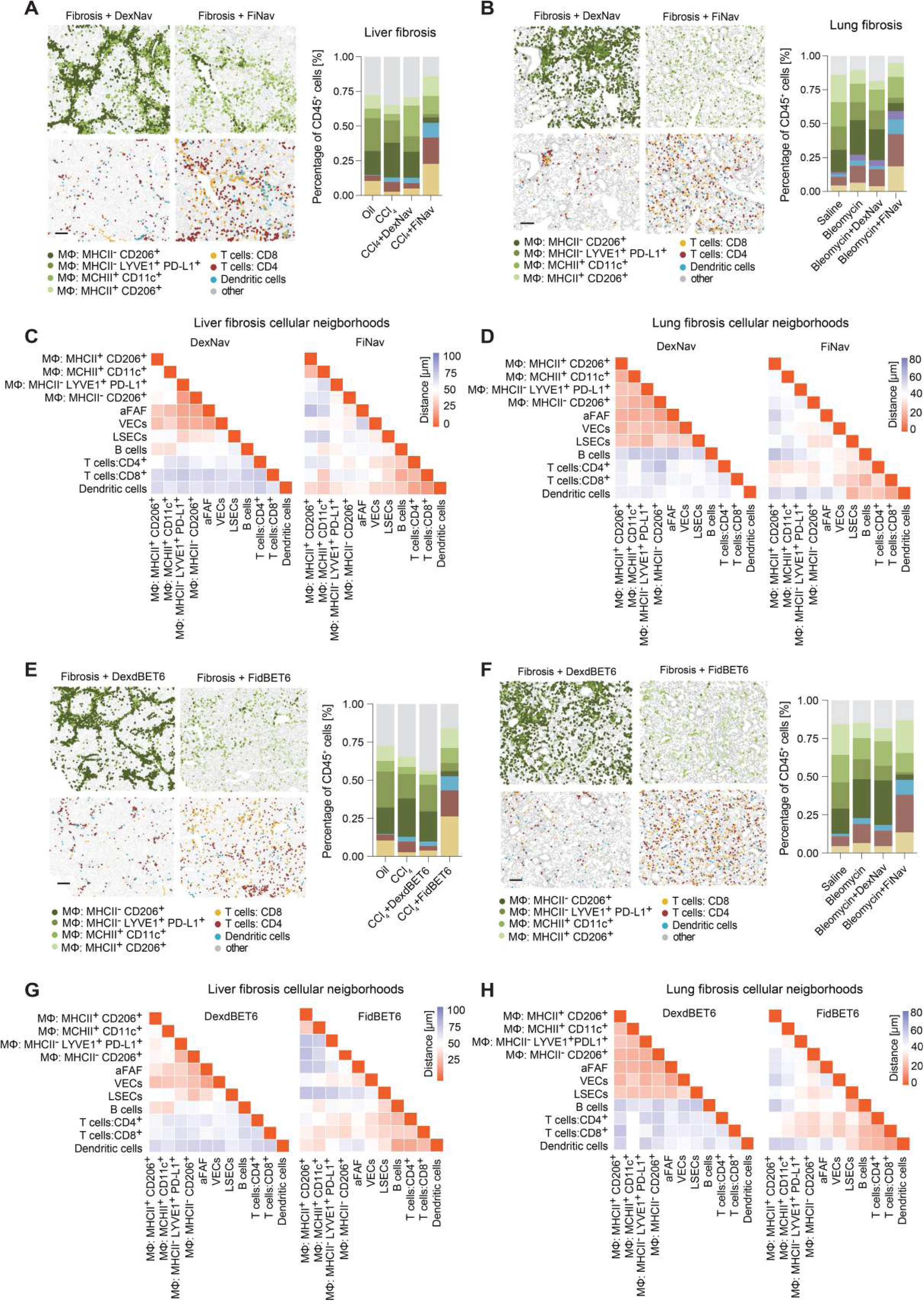
Spatial characterization of the immune microenvironment remodeling in fibrotic tissues upon SMNP treatment. (Related to Figure 4). (**A-B**) (Left) Representative images of mIF analysis mapping immune and stromal populations in murine fibrotic livers (A) and lungs (B) treated for four weeks with DexNav or FiNav nanoparticles. Scale bars, 250 μm. (Right) Quantification of the indicated immune and stomal populations. (**C-D**) Cellular neighborhood analysis measuring the distance between the indicated immune and stromal populations in murine fibrotic livers (C) and lungs (D) treated for four weeks with DexNav or FiNav. (**E-F**) (Left) Representative images of mIF analysis mapping immune and stromal populations in murine fibrotic livers (E) and lungs (F) treated for four weeks with DexdBET6 or FidBET6. Scale bars, 100 μm. (Right) Quantification of the indicated immune and stomal populations. (**G-H**) Cellular neighborhood analysis measuring the distance between the indicated immune and stromal populations in murine fibrotic livers (C) and lungs (D) treated for four weeks with DexdBET6 or FidBET6.

**Figure S10:**
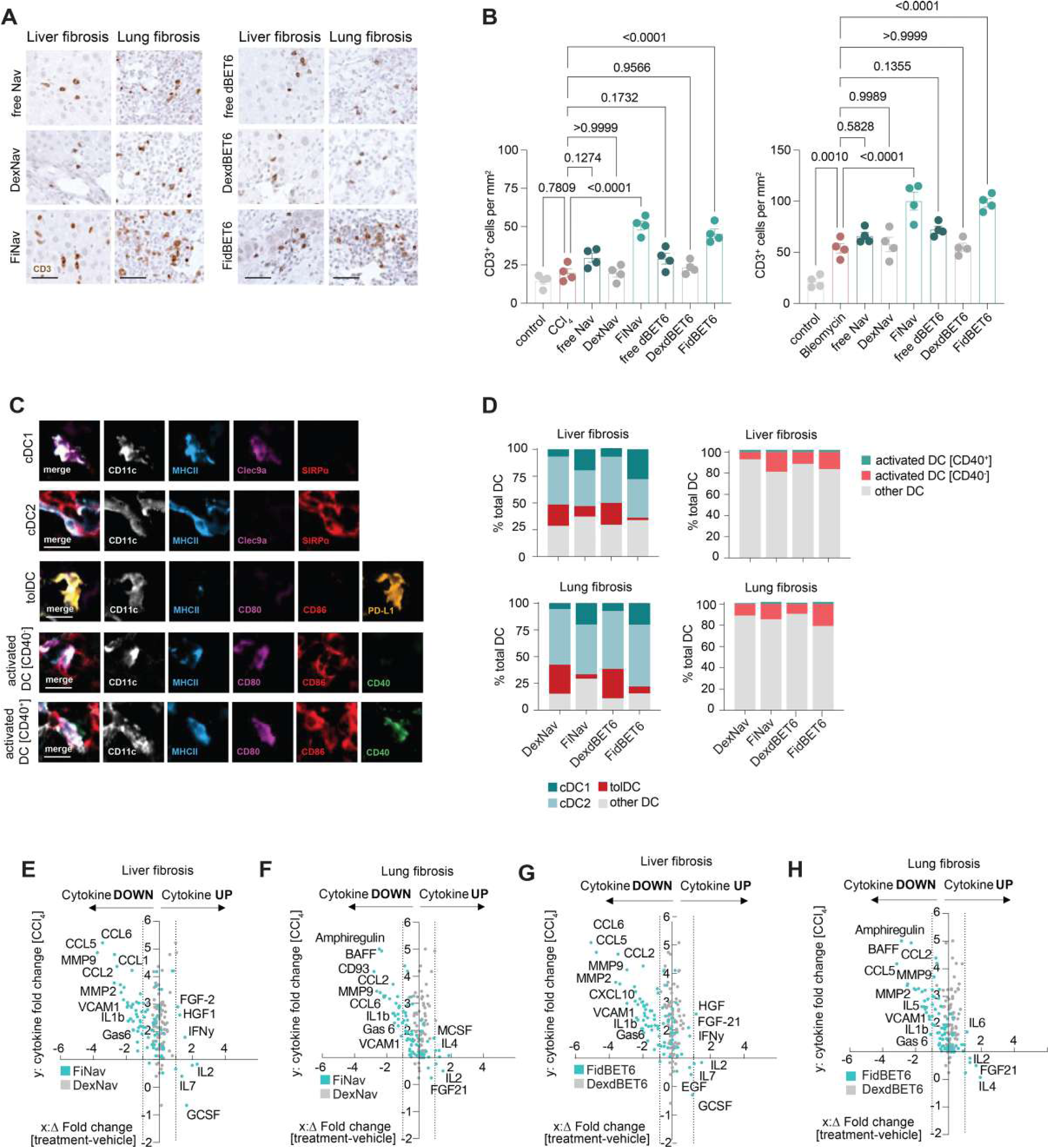
Impact of SMNPs on immune cells and cytokines in fibrotic, immunosuppressive niches. (Related to Figure 4). (**A**) Representative immunohistochemistry analysis of CD3^+^ T cells in fibrotic liver (top) and lungs treated with free navitoclax, free dBET6, DexNav, DexdBET6, FiNav and FidBET6 for 4 (liver) or 3 (lung) weeks. Scale bars, 75 μm. (**B**) Quantification from A in fibrotic livers (left) and lungs (right) (4 mice/group). (**C-D**) Representative mIF pictures of conventional DC 1 (cDC1), cDC2, tolerogenic DC (tolDC), and CD40^+^ or CD40^-^ activated DCs in murine fibrotic livers (C) and quantification in murine fibrotic livers and lungs treated for four weeks with DexNav, FiNav, DexdBET6, or FidBET6 nanoparticles (D). (**E-F**) Differential protein expression of cytokines from whole-tissue lysates of murine fibrotic livers (E) or lung (F) after 4 (liver) or 3 (lung) weeks of DexNav or FiNav nanoparticle treatment. (**G-H**) Differential protein expression of cytokines from whole-tissue lysates of murine fibrotic livers (G) or lung (H) after 4 (liver) or 3 (lung) weeks of DexdBET6 or FidBET6 nanoparticle treatment. Each dot represents one mouse (B). Data are presented as mean ± s.e.m. The one-way ANOVA with Sidak’s multiple comparisons test was used for (B).

**Figure S11:**
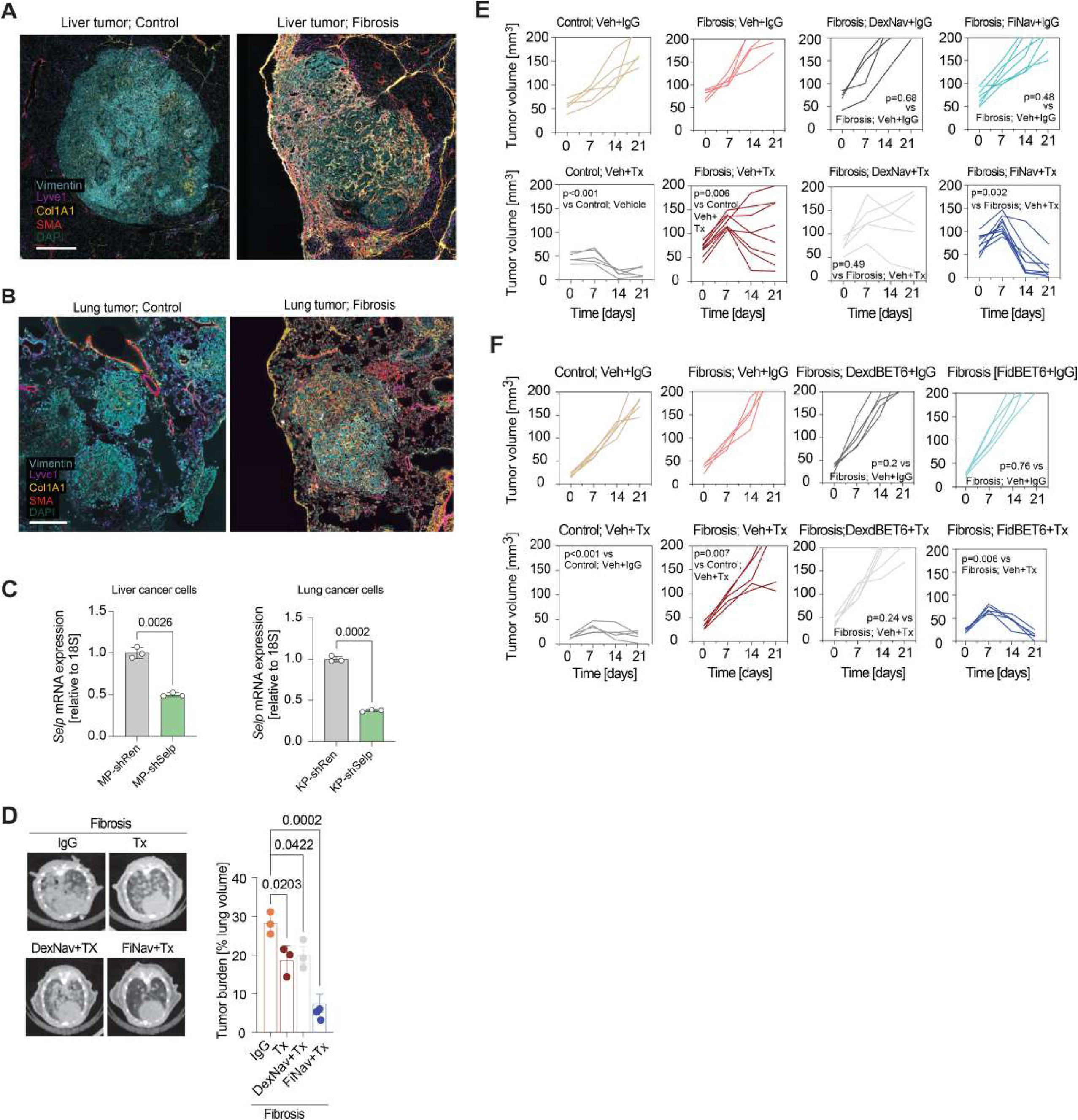
Impact of SMNPs on ICB responses in mice with fibrotic liver and lung tumors. (Related to Figure 5). (**A**-**B**) mIF analysis of vimentin, Lyve1, Collagen type I (Col1A1), and smooth muscle actin (SMA) expression on fibrotic and non-fibrotic murine liver (A) and lung tumors (B). Scale bars, 250 μm. (**C**) Quantitative reverse transcription polymerase chain reaction (RT-PCR) analysis of *Selp* mRNA isolated from MP liver cancer cells (left) and KP lung cancer cells (right). 18S was used as the internal amplification control. (**D**) (Left) Representative image of microCT scan at d14 of murine fibrotic lung tumors treated with DexNav or FiNav nanoparticles in combination with lung first-line Tx. (Right) Quantification of microCT scans (n=3 mice/group). (**E**) Quantification of tumor volume change over time in murine fibrotic liver tumors treated with DexNav or FiNav nanoparticles in combination with liver first-line Tx (5-10 mice/group). (**F**) Quantification of tumor volume change over time in murine fibrotic liver tumors treated with DexdBET6 or FidBET6 nanoparticles in combination with liver first-line Tx (5-10 mice/group). Each dot represents one biological replicate (C), one mouse (D). Each line (E-F) represents one mouse. Data are presented as mean ± s.e.m. Unpaired t-test was used for (C). Statistical tests were the one-way ANOVA with Sidak’s multiple comparisons test was used for (D), and the Mann-Whitney test or Welch’s t-test at endpoint (21 days) was used for (E-F).

**Figure S12:**
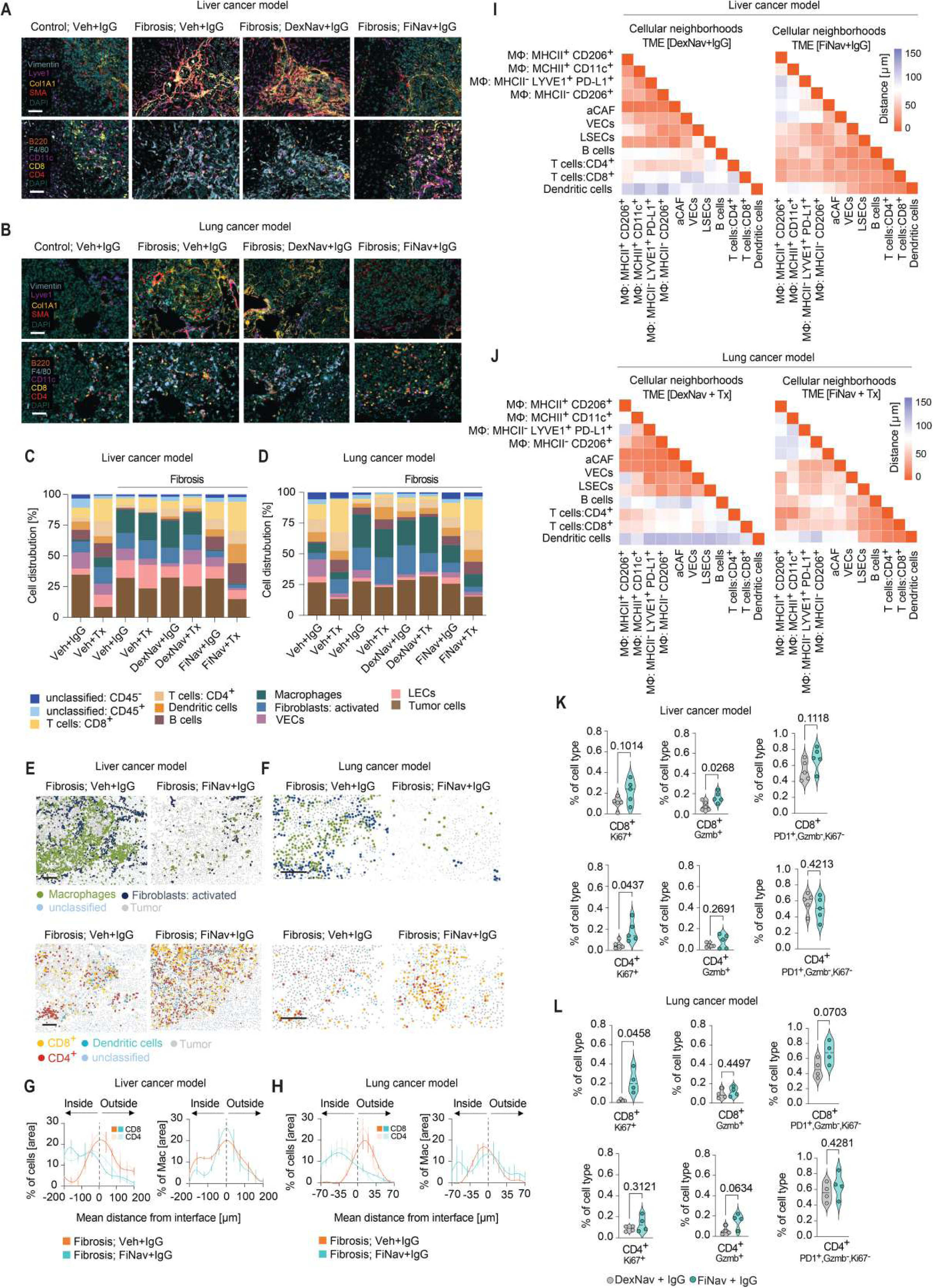
SMNP-mediated niche remodeling in fibrotic liver and lung tumors. (Related to Figure 5). (**A-B**) mIF analysis of stromal (Vimentin, Lyve1, Col1a1, SMA) and immune (B220, F4/80, Cd11c, CD8, CD4) markers in murine fibrotic liver (A) or lung (B) tumors treated with DexNav or FiNav nanoparticles in combination with IgG controls. Scale bars, 100 μm. (**C-D**) Proportion of immune and stromal cells in murine fibrotic liver (C) or lung (D) tumors treated with DexNav or FiNav nanoparticles in combination with first-line Tx. (**E-F**) Representative images of mIF analysis mapping different immune (top) and stromal (bottom) populations in murine fibrotic liver (E) or lung (F) tumors treated with FiNav nanoparticles in combination with IgG controls. Scale bars, 100 μm. (**G-H**) Proportion of CD8^+^, CD4^+^ T cells (left) and macrophages (right) present inside or outside the tumor region in murine fibrotic liver (G) or lung (H) tumors treated with FiNav nanoparticles in combination with IgG controls. (**I-J**) Cellular neighborhood analysis measuring the distance between the different immune and stromal populations in murine fibrotic liver (I) or lung (J) tumors treated with DexNav or FiNav nanoparticles in combination with first-line Tx. (**K-L**) Quantification of active/cytotoxic/exhausted CD8^+^ and CD4^+^ T cells in murine fibrotic liver (K) or lung (L) tumors treated with DexNav or FiNav nanoparticles in combination with IgG controls (4-5 mice/group). Each line or dot represents one mouse. Data are presented as mean ± s.e.m. Statistical tests were Mann-Whitney test or Welch’s t-test (K, L).

**Figure S13:**
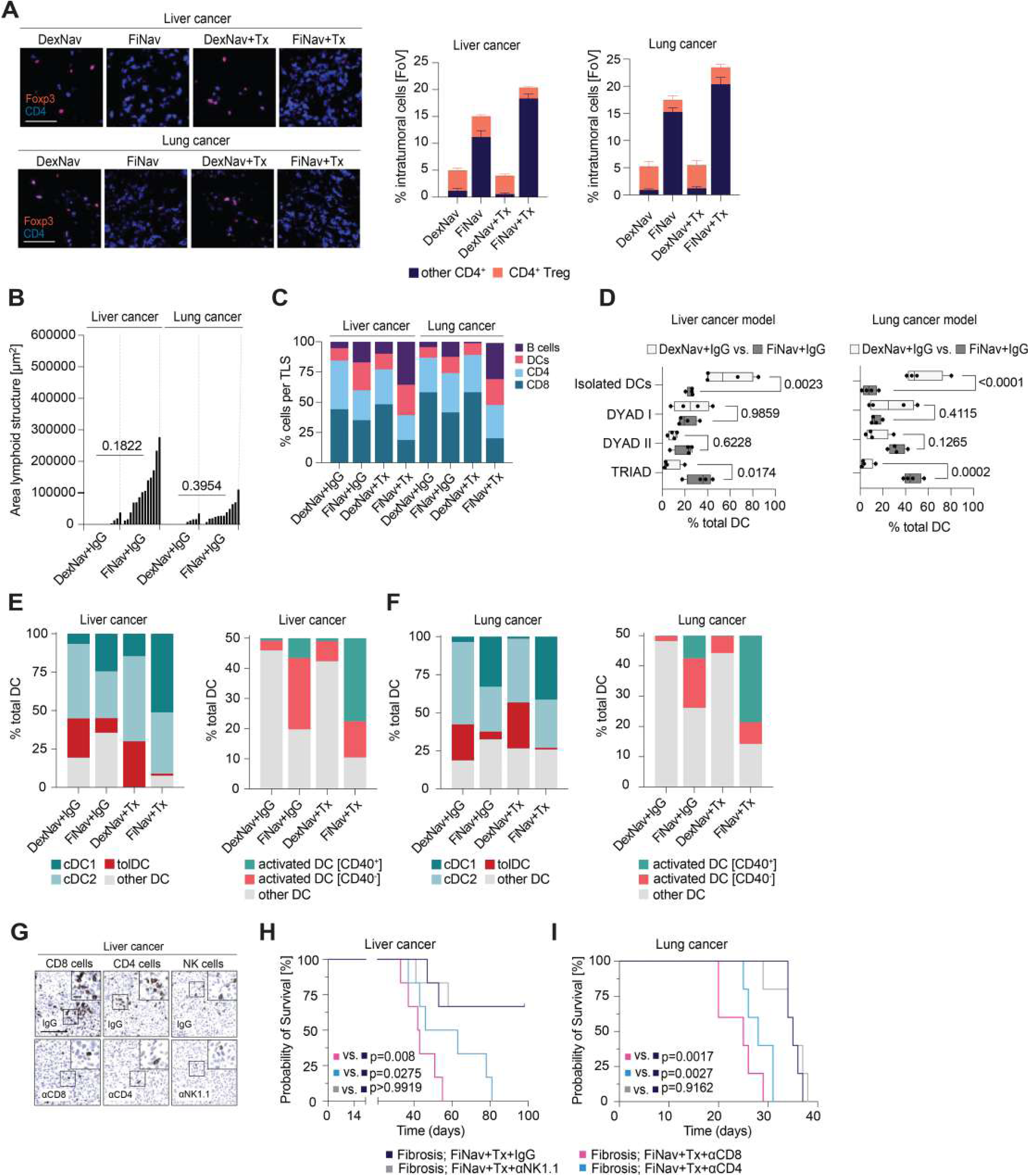
Impact of SMNPs on lymphoid aggregates in the tumor microenvironment and dependence on CD8^+^ T cells. (Related to Figure 5). (**A**) Representative IF images showing Foxp3^+^ regulatory T cells (Treg) and CD4^+^ T changes during SMNP treatments. Quantification of CD4^+^ and Treg in murine fibrotic liver (left) or lung (right) tumors treated with DexNav or FiNav nanoparticles in combination with first-line Tx. (**B**) Quantification of lymphoid aggregates in murine fibrotic liver and lung cancer treated with DexNav or FiNav nanoparticles in combination with IgG controls, related to figure 5M. (**C**) Proportions of B cells, dendritic cells, CD8^+^ T cells, and CD4^+^ T cells in lymphoid aggregates in murine fibrotic liver and lung cancer treated with DexNav or FiNav in combination with liver/lung first-line Tx (or IgG controls) (4 mice/group). (**D**) Quantification of isolated DCs, DC::CD4 dyads (DYAD I), DC:CD8 dyads (DYAD II), and DC::CD4::CD8 triads (TRIAD) in murine fibrotic liver (left) and lung (right) cancer treated with DexNav or FiNav in combination with IgG controls. (**E-F**) Quantification of (left) conventional DC 1 (cDC1), cDC2, and tolerogenic DC (tolDC) or (right) CD40^+^ and CD40^-^ activated DCs in murine fibrotic livers (E) and lung (F) cancer treated for four weeks with DexNav, FiNav, DexdBET6, or FidBET6. (**G**) Representative immunohistochemistry images of CD8, CD4, natural killer (NK) cells upon CD8, CD4, or NK cell depletion in murine fibrotic liver cancer. (**H-I**) Kaplan-Meier analysis comparing overall survival of mice with fibrotic liver (H) and lung (I) tumors treated with FiNav nanoparticles in combination with first-line Tx and depletion antibodies for either CD8^+^ T cells, CD4^+^ T cells, NK1.1^+^ natural killer cells, or IgG controls. Each line or dot represents one mouse. Data are presented as mean ± s.e.m. Statistical tests were the Mann-Whitney test for (B), the one-way ANOVA with Sidak’s multiple comparisons test for (D), and the log-rank Mantel-Cox test for (H, I).

**Figure S14:**
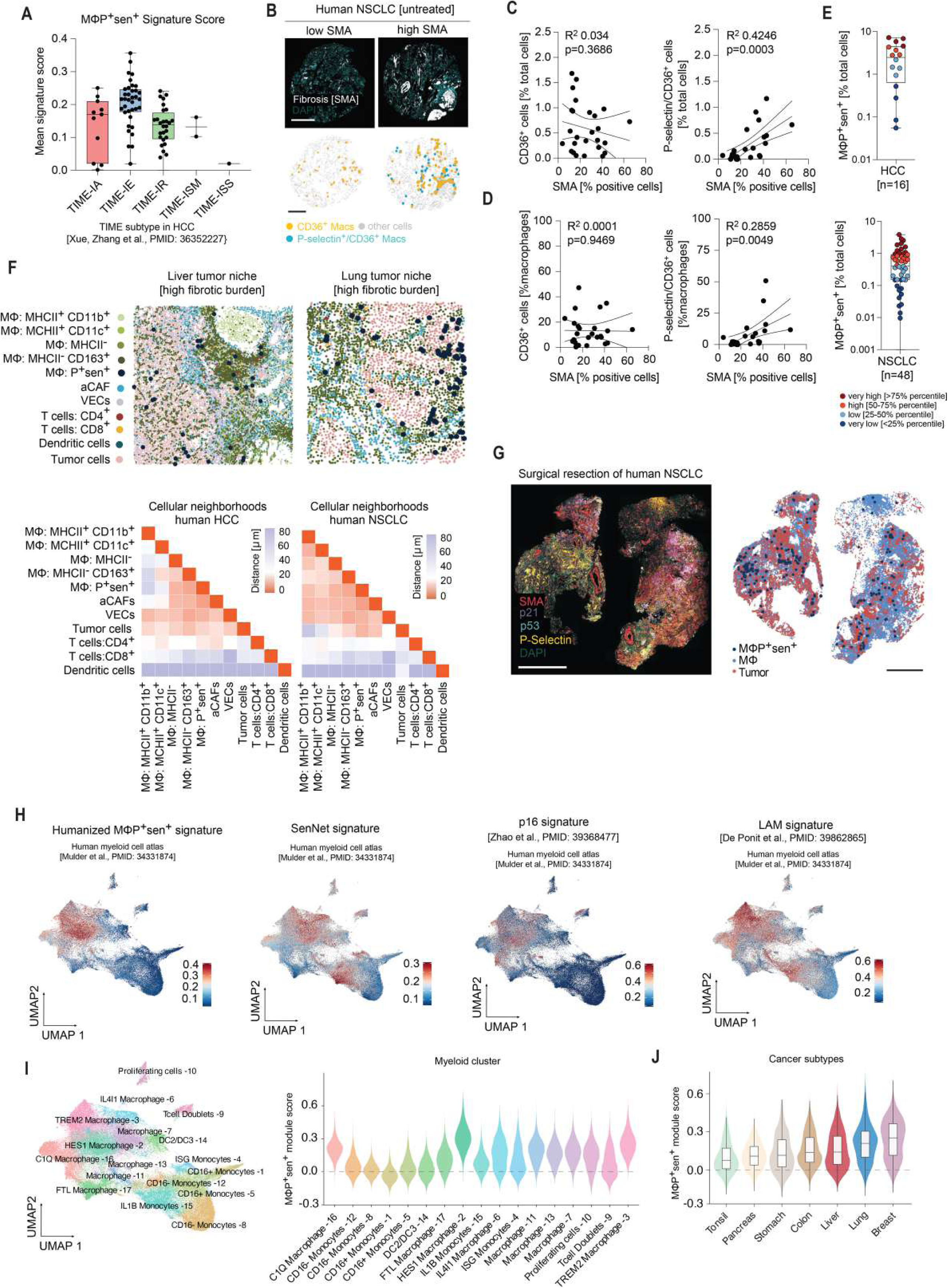
Characterization of P^+^sen^+^ macrophages in human tumors. (Related to Figure 6). (**A**) Signature score of P^+^sen^+^ macrophages across previously defined tumor immune microenvironment (TIME) classes of human HCC: Immune active (IA; n=11), Immune excluded (IE; n=36), Immune resident (IR, n=28), Immune suppressive myeloid (ISM; n=2) and Immune suppressive stromal (ISS; n=1). (**B**) Representative core image of mIF analysis mapping CD36^+^ and P-selectin^+^/CD36^+^ macrophages in human NSCLC with a low or high fibrotic burden. Fibrotic burden has been quantified by a-smooth muscle actin (aSMA) expression. Scale bars, 250 μm. (**C**) Linear regression analyses between (left) CD36^+^ cells (normalized to total cell number) and activated fibroblasts (aSMA) and (right) P-selectin^+^CD36^+^ cells (normalized to total cell number) and activated fibroblasts (aSMA) in human NSCLC tissues (26 patients) (**D**) Linear regression analyses between (left) CD36^+^ cells (normalized to total macrophages) and activated fibroblasts (SMA) and (right) P-selectin^+^CD36^+^ cells (normalized to total macrophages) and activated fibroblasts (SMA) in human NSCLC tissues (26 patients). (**E**) IF quantification of P^+^sen^+^ macrophages in human HCC (top) and NSCLC (bottom) related to Fig. 6G-H. (**F**) (Top) Representative image of mIF analysis mapping different immune and stromal populations in human liver (left) and lung (right) tumor niche with high fibrotic burden (determined by aSMA level). (Bottom) Cellular neighborhood analysis measuring the distance between the different immune and stromal populations in human HCC (left) or NSCLC (right) tumors (analyzed in high fibrotic HCC (n=3)/NSCLC(n=3) samples). Scale bars, 150 μm. (**G**) Representative image of multiplex analysis mapping stromal, tumor, and macrophage populations in surgically resected human NSCLC tumor. Scale bars, 500 μm. (**H**) Humanized P^+^sen^+^ macrophage, SenNet, p16, and LAM signature in cross-tissue scRNA datasets of human myeloid cells in health and disease. (**I**) (Left) Cell type annotation and clustering as annotated by Mulder et al. (PMID: 34331874) and (right) P^+^sen^+^ macrophage signature score in each cluster. (**J**) P^+^sen^+^ macrophage signature score across cancer types. Each dot represents one patient (A, C-E). Data are presented as mean ± s.e.m. Simple linear regression was used for (C-D).

